# Spatial and seasonal biodiversity variation in a large Mediterranean lagoon using environmental DNA metabarcoding through sponge tissue collection

**DOI:** 10.1101/2024.10.17.618912

**Authors:** Andrea Corral-Lou, Ana Ramón-Laca, Lourdes Alcaraz, Robert Cassidy, Sergi Taboada, Stefano Mariani, Borja Mercado, Martin Vicente, Ángel Pérez-Ruzafa, Ramón Gallego, Ana Riesgo

## Abstract

Ecosystem monitoring is a fundamental tool to avert biodiversity loss, gathering valuable information that can be used to develop conservation policies, evaluating management outcomes, and guiding science-based decision-making. The Mar Menor costal lagoon (South-East of Spain) has experienced episodes of eutrophication due to intensive agriculture and other human activities, causing mass mortalities of marine fauna. In this scenario, biodiversity monitoring is crucial to evaluate the status of fauna and flora and take appropriate measures. Here, our main objective was to assess faunal composition and its spatial and temporal structure associated with the pillars used to support the built recreational well-being facilities along the Mar Menor. We capitalised on the many sea sponges that settle on these structures by collecting tissue samples for subsequent extraction of filtered environmental DNA (i.e. ‘natural sampler DNA’, nsDNA) in northern and southern areas of the lagoon, both in spring and summer. We metabarcoded all samples at the cytochrome oxidase subunit I gene (*COI*), and reliably identified 76 taxa belonging to nine different phyla, with annelids, poriferans, molluscs and cnidarians being the richest groups. We detected emblematic species of threatened status, such as pipefishes (genus *Syngnathus*) and the piddock clam, *Pholas dactylus*, and others known to become invasive, such as the ascidians *Styela canopus* and *Botrylloides niger*, the barnacle *Amphibalanus amphitrite*, and the polychaetes *Branchiomma bairdi* and *Branchiomma boholense*. The use of common and widely distributed sponges as natural eDNA samplers allowed us to characterise both spatial and temporal diversity, further emphasizing the importance of this low-cost approach to monitoring biological communities in shallow coastal ecosystems.

## Introduction

Coastal lagoons are unique because they form a continuum between continental and marine aquatic ecosystems; they constitute sheltered habitats, variably isolated from the open sea, and with unique heterogeneity features created by freshwater inputs, salinity anomalies, and distinctive biogeochemical cycles, which boost their biological productivity (Pérez-Ruzafa et al. 2011). Lagoons habitats are diverse and frequent along the continental margins, occupying only 13% of the world’s coastline (Kjerfve, 1994). However, as a result of the interplay between sediment accumulation from rivers, coastal hydrology, and sheet geomorphology, they tend to be particularly vulnerable to pollution, modifications, and even destruction, as a result of human commercial, industrial, and recreational activities, highlight the importance of their conservation. The Mar Menor lagoon, located in South-East of Spain, is one of the largest coastal lagoons in the Western Mediterranean, with a surface area of 136.1 km^2^ and a maximum depth of 7 meters (García-Oliva et al., 2018). The physical and geographic features of the Mar Menor support its significant ecological role, hosting a unique community of organisms that are specially adapted to live under variable salinity conditions (Pérez-Ruzafa et al., 2005), and which contribute to the rich biodiversity of the area (Alongi, 1998, Kennish & Paerl, 2010; Clark, 1998; Vasconcelos et al., 2011; Yañez-Arancibia & Nugent, 1977; Pérez-Ruzafa et al., 2019a). Apart from its ecological importance, the Mar Menor also provides other ecosystem services for humans, such as tourism and the provision of resources for agriculture (Velasco et al., 2018), which in turn has resulted in frequent human disturbances. These activities put at risk the lagoon’s ecological integrity and the goods and services it provides, such as water and bathing quality, fishing yields and the capacity to withstand eutrophication or retain heavy metals in its sediments (Pérez-Ruzafa et al., 1991, 2023; Marcos et al., 2015; Vázquez-Luis et al., 2017; Fernández-Alías et al., 2022). One such disturbance is an increase in the volume of water exchanged between the Mediterranean and the Mar Menor in recent decades, which has been linked to the deepening of the El Estacio channel for the construction of a port and make it navigable (Arévalo, 1988; Pérez-Ruzafa et al., 2005). Additionally, there has been an increase in nitrate concentrations due to the intensification of agriculture in the area over the last decade (Pérez-Ruzafa & Marcos, 2004), which has caused the Mar Menor to shift from moderately oligotrophic to severely eutrophic status (Pérez Ruzafa et al., 2002, 2019; Sadonnini et al. 2021). These two factors among others like the tourism pressure (Conesa & Jiménez-Cárceles 2007), have contributed to recent changes in the biological communities of the lagoon by facilitating the arrival of new species with the potential to alter ecosystem functioning (Pérez Ruzafa & Aragón, 2003; Sandonnini et al., 2021), as well as by contributing to episodes of mass mortality throughout the lagoon and anoxia in deeper areas, affecting the general biodiversity and the viability of some emblematic species of the area (Belando et al., 2017; Giménez-Casalduero et al., 2020; Sandonnini et al., 2021; Fernández-Alías et al., 2022).

In recent years, the environmental DNA (eDNA) approach, based on the screening of the genetic material released into the environment by organisms through the shedding of cells and tissues (Taberlet et al., 2012), has been widely used to monitor biodiversity (Goldberg et al.,2015; Valentini et al., 2016; Cilleros et al., 2019; Sales et al., 2021). The use of eDNA as a monitoring tool has certain advantages over traditional methods and is particularly effective at generating an initial comprehensive assessment of the biodiversity in an ecosystem. Notably, this technique does not require the direct collection of specimens, which can compromise the viability of biological communities, and is a particularly sensitive issue when a taxon is threatened (Piaggio, 2021; Sahu et al., 2023). Furthermore, traditional monitoring across wide taxonomic ranges also needs expert taxonomists to morphologically identify the organisms under study, resulting in elevated costs (Snyder, 2003; Roy et al., 2018; Bylemans et al., 2019; Goutte et al., 2020). In contrast, the non-invasive technique of eDNA monitoring, has demonstrated a higher sensitivity to detect species in marine ecosystems than traditional methods (Hinlo et al., 2017; Sigsgaard et al., 2020; Fraija-Fernández et al., 2020; Heino et al., 2011), and a great capacity to determine the composition of biodiversity at different spatial and temporal scales (Jeunen et al., 2018; Berry et al., 2019; Jeunen et al., 2020; West et al., 2020).

Typically, eDNA is collected by filtering a volume of water sample through a filter made from a material such as cellulose nitrate, isopore membrane, glass fibre or isopore polycarbonate (Rees et al., 2014). Alternatively, the so-called “natural samplers”, which are typically organisms that exhibit filter-feeding behaviour, provide a strong alternative to water or sediment collection as a method of obtaining eDNA (nsDNA, see Mariani et al., 2019; Weber et al., 2023), as long as they are abundant and resilient. In particular, sea sponges have been shown to be extraordinary natural samplers of eDNA, having previously enabled the detection of a high proportion of the vertebrate communities in different marine environments (Mariani et al. 2019, Turon et al. 2020, Brodnicke et al. 2023, Neave et al. 2023, Jeunen et al. 2023, Cai et al. 2024, Gallego et al., 2024).The eDNA amplified from sponge tissues is also detectable for longer periods of time than that collected from seawater column samples (Cai et al., 2022), and has been found to outperform biodiversity estimates from eDNA collected from seawater, traditional catchment methods, and acoustic biomonitoring (Cai et al., 2024). The use of sponges as natural samplers therefore yields great advantages, simplifying the data collection workflow and providing an extremely powerful and cost-effective alternative for biodiversity monitoring, particularly in inaccessible ecosystems (Mariani et al., 2019; Turon et al., 2020; Brodnicke et al., 2023; Harper et al., 2023; Neave et al., 2023; Jeunen et al., 2023; Cai et al., 2024; Gallego et al., 2024). Since the discovery of sponges’ utility as effective natural samplers of eDNA, various representatives of the class Demospongiae and Hexactinellida have been used as natural eDNA samplers (e.g., Turon et al., 2020; Cai et al., 2022; Harper et al., 2023, Brodnicke et al. 2023; Gallego et al., 2024). Importantly, different species of sponges have been found to exhibit different filtering characteristics and microbial composition and abundance in their bodies, features that have been found to be correlated to variability in the efficiency of eDNA accumulation, and which suggests that some sponges are better natural samplers than others (Brodnicke et al., 2023; Cai et al., 2023, Neave et al. 2023, Gallego et al. 2024).

In the Mar Menor costal lagoon, there is a variety of sponge species of different genera (Pérez-Ruzafa, 1989), including *Haliclona mediterranea* Griessinger, 1971, *Haliclona elegans* (Lendenfeld, 1887), *Haliclona oculata* (Linnaeus, 1759), *Halichondria semitubulosa* (Lamarck, 1814), *Suberites massa* Nardo, 1847, *Dictyonella incisa* (Schmidt, 1880), *Sycon raphanus* Schmidt, 1862, and species of recent colonization as *Dysidea fragilis* (Montagu, 1814), and *Cliona celata* Grant, 1826. However, no previous studies have assessed the ability of any of these species to act as natural eDNA samplers. Furthermore, the taxonomy of several species of the genus *Haliclona* reported in the Mar Menor is believed to be uncertain (De Weerdt, 2002). *Haliclona* sp. are also classified as Low Microbial Abundance (LMA) sponges in terms of their microbiome composition, making them good candidates as natural samplers of eDNA (Moitinho-Silva et al. 2017), as shown by earlier nsDNA studies using *H. scotti* (Kirkpatrick, 1907) or *Haliclona toxotes* (Hentschel, 1912) (Turon et al., 2020; Jeunen et al., 2023). In addition, these species are widespread throughout the Mediterranean and in some areas of the eastern Atlantic, which could facilitate their large scale inclusion in next-generation biomonitoring strategies.

The sponges, specifically the species *Haliclona mediterranea*, are one of the most dominant species in the “balnearios” (piers fitted out as bathing areas) of the Mar Menor lagoon along with others filter-feeders (Perez-Ruzafa et al., 2019, 2020). These small piers across the lagoon, constructed with wooden or concrete piling since 1987 to facilitate bathing, have provided structural shaded habitats that favour the settlement and development of sciaphilic assemblages and enhance connectivity (Pérez-Ruzafa et al., 2005; Marchini et al., 2004; Firth et al., 2016). However, their presence has also been considered responsible for the spread of both autochthonous and allochthones species by providing suitable stepping stone habitats, allowing the colonisation of previously inaccessible areas, and overall favouring the establishment of fast colonisers in these areas (Glasby et al. 2007; Sanabria-Fernandez et al., 2018). Here, we used biopsies of *Haliclona* sponges settled on these artificial structures as eDNA capsules to assess the biodiversity of sciaphilic communitiesassociated with these small piers, and tracked seasonal changes in species composition through metabarcoding of eDNA based on the *COI* marker.

## Materials & Methods

### Sampling site

The investigated areas are located in the Mar Menor lagoon, in the southeastern region of the Iberian Peninsula (Figure 1). A total of four sample sites were studied (Table 1; Figure 1), of which two were located on the pillars under two different docks and piers with sciaphilic communities at La Ribera (37.805674, -0.797644 and 37.805478, -0.797052) and two in the same type of structures at Los Urrutias (37.695094, -0.838869 and 37.684317, -0.830104). All sampling was performed at depths of less than two meters, with sampling carried out in both spring and summer of 2022 to account for seasonal differences. Approximately 5 cm^3^ of tissue from four sponge specimens (*Haliclona* spp.) were sampled at each sample site and season, yielding a total of 16 specimens. Each sponge fragment was collected in an individual, sterile, DNAase-free tube and stored in absolute ethanol at -20°C overnight; then, the ethanol was replaced twice and stored at -20°C until further processing.

**Figure 1.**
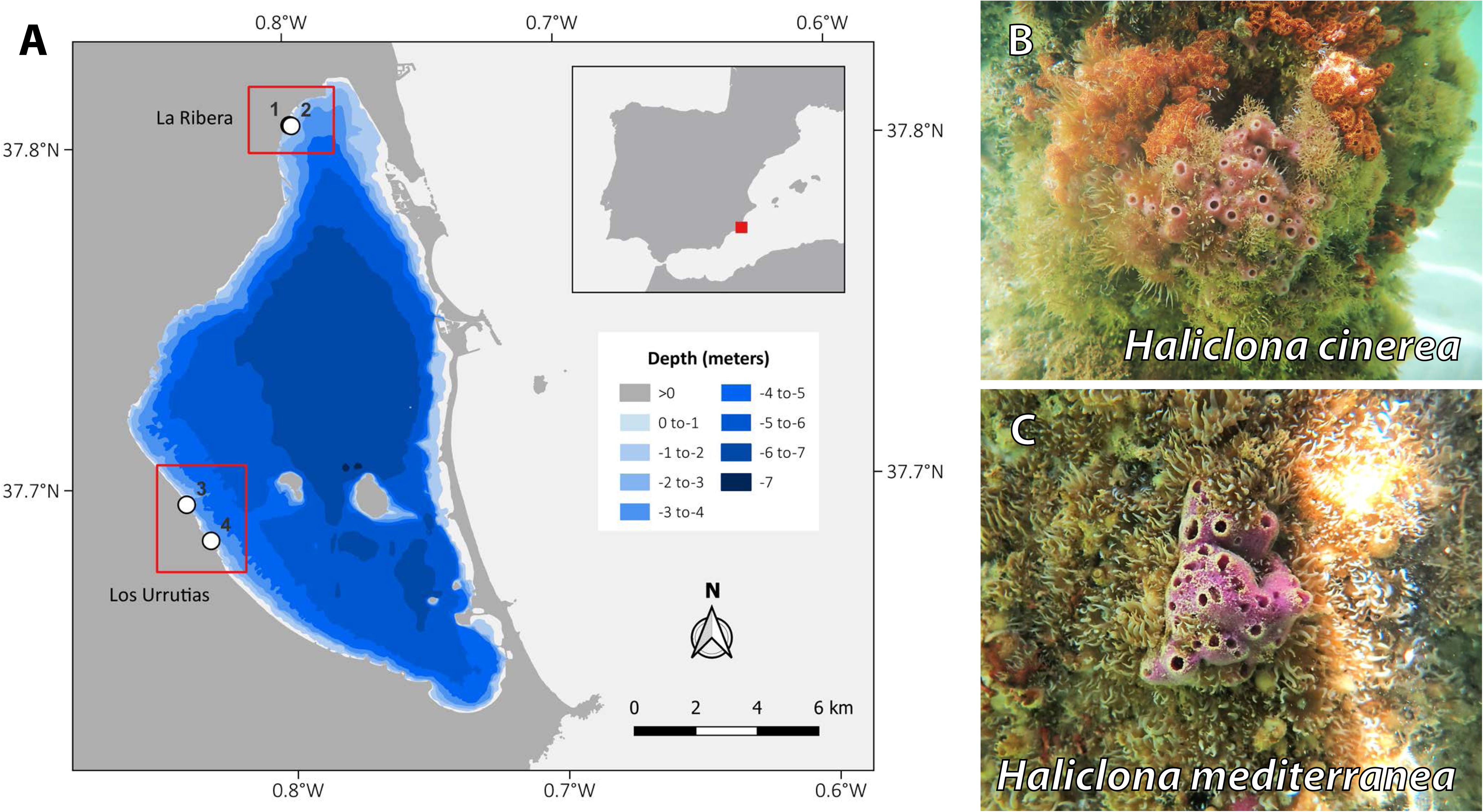
A) Sampling sites included in this study. The blue shading indicates the different depths (more details in the legend). Numbers correspond to those in Table 1. B) *Haliclona cinereal* (in dark pink), the dominant host sponge in Los Urrutias, surrounded by the invasive ascidian *Botrylloides niger* (in orange). C. *Haliclona mediterranea* (in purple), the dominant host sponge in La Ribera, flanked by the anthozoan *Exaiptasia diaphana* (in yellow).

**Table 1.** Summary of the samples collected and analyzed in this study

| Locality | Site | Sponge | Season | PCR | Total |
| --- | --- | --- | --- | --- | --- |
| La Ribera (Ri) | 2 sites (1 and 2) | 4 sponges (1, 2, 3 and 4) | Spring and Summer (Sp and Sum) | 3 replicates (rep1, rep2 and rep3) | 48 |
| Los Urrutias (Ur) | 2 sites (3 and 4) | 4 sponges (1, 2, 3 and 4) | Spring and Summer (Sp and Sum) | 3 replicates (rep1, rep2 and rep3) | 48 |
| Positive control | Tissue of the freshwater fishes *Pangasionodon hypophthalmus* | - | - | 3 replicates (rep1, rep2 and rep3) | 3 |

### Environmental DNA sampling, extraction, library preparation and sequencing

The DNA from a fragment (approximately 0.5 cm^3^) of each sponge individual was extracted. For each individual, after ethanol removal and blotting, DNA was extracted using the Speedtools Tissue DNA Extraction Kit (Biotools ® B&M Labs SA, Spain) following the manufacturer’s protocol but with an overnight lysis incubation. After extraction, DNA was eluted in two steps to a final volume of 150 µl to maximize DNA yield. Metazoan taxa were targeted for analysis by amplifying a 313-bp segment of the mitochondrial *cytochrome* c *oxidase subunit I* gene (*COI*) using the forward primer mlCOIintF-XT (5’-GGW ACW RGW TGR ACW ITI TAY CCY CC-3′; Wangensteen et al., 2018, a modified version of mlCOIintF of Leray et al., 2013) and the reverse primer jgHCO2198 (5’-TAI ACY TCI GGR TGI CCR AAR AAY CA-3’; Geller et al., 2013). A positive control, DNA from *Pangasionodon hypophthalmus*, a freshwater fish not found in the Iberian Peninsula, was included with the samples. The eDNA samples were amplified in triplicate in 20-μl reactions containing: 10 μl MyFi (Meridian Bioscience), 0.16μl BSA (Thermo Fisher Scientific), 1μl of the primer mix (forward and reverse at a concentration of 10 µM) (Thermo Fisher Scientific), 2 μl of DNA template (≤20 ng µl^-1^) and 6.84 μl of molecular grade water. Tagged primers consisted of an 8-bp dual barcode and between 1-4 random nucleotides added to improve sequence diversity for Illumina processing, ensuring optimal nucleotide diversity at each sequencing cycle. PCR cycle conditions were set to the following parameters: initial denaturation step at 95°C for 10 min, followed by 45 cycles of 95°C for 45 s, 45°C for 45 s and 72°C for 45 s, with a final extension cycle of 5 min. The concentrations of PCR products were measured using a Qubit dsDNA HS Assay kit (Invitrogen). Amplicons were pooled to obtain ≥ 3 µg of total DNA, aiming for 100 µl when possible, by either evaporating the resulting volume to 100 µl if necessary or correcting for the actual volume proportion in the next step. Size selection for fragments of the desired size was performed in two washes using Ampure XP beads, first at 0.5× to remove fragments longer than 400 bp, then at 0.8× to keep fragments longer than 300-bp. The libraries were imaged on a Tape Station 4200 (Agilent) using an Agilent high sensitivity D1000 tape station kit to check the purity and average base pair length. Each library was then ligated using unique adapters, including the i7 and i5 library barcodes of NEXTFLEX® Rapid DNA-Seq Kit 2.0 for Illumina (PerkinElmer), following manufacturer’s instructions with 8 PCR cycles using 500 ng of library, and imaged again on the TapeStation to check for an increase in average base-pair length. Libraries were quantified using the Quant-iT dsDNA HS assay kit with a Qubit® 2.0 Fluorometer (Life Technologies) and pooled at equimolar ratios into a final mix. The single resulting library was paired-end sequenced at 12.5 pM with 10% PhiX on an Illumina MiSeq using a V3-600 sequencing kit at the Genomics Division of the Universidad Complutense de Madrid producing 300-bp paired-end reads.

### Bioinformatic processing

Raw reads were initially separated into three paired-end libraries by Illumina’s proprietary software, and further demultiplexed into the starting 99 uniquely barcoded samples consisting of three positive controls and three replicates of the 32 unique biological samples. We performed this operation using a custom pipeline combining bash scripts and the cutadapt v4.2 program (Martin, 2011), which also carried out primer removal and determined the direction in which amplicons were sequenced (available on https://github.com/ramongallego/Mar_Menor).

Amplicons were trimmed and quality filtered using FilterAndTrim from the DADA2 R package (Callahan et al. 2016) with the following parameters: *truncLen=c(220,160)*, *maxN=0* and *maxEE=c(2,2)*. Through the different functions of DADA2, we also estimated and corrected the errors, performed the dereplication step and merged the paired sequence, obtaining final dataset of the correct Amplicon Sequence Variants (ASVs). Resulting ASVs were further clustered into OTUs using a method based on single-linkage-clustering (SWARM) (Mahé et al., 2014) with a distance of two steps within each sample, which was chosen due to its good performance compared to other methods to reduce sequencing errors and PCR artifacts (Kopylova et al., 2016). The removal of chimeras was carried out using VSEARCH v2.21.1 (Rognes et al., 2016).

### Taxonomic assignment

Taxonomic assignment was performed using the BLAST tool (Altschul et al., 1997) to align the sequences with the *nr* database from the National Center for Biotechnology Information (NCBI) GenBank nucleotide database with the following parameters: *-perc_identity 75 -word_size 30 -evalue 1e-30 -max-target-seqs 50 -culling_limit 5.* Once all matching sequences were obtained, records that had a percent similarity of less than 90% and/or an alignment coverage of less than 90% were removed. Matches in which the organism was not identified, and sequences were uploaded as “environmental samples” were also removed. Final taxonomical identification was performed using the function *condenseTaxa* from the R package taxonomizer. The r script parsing arguments and defining all necessary thresholds can be found in the supplementary material (available on https://github.com/ramongallego/Mar_Menor).

### Barcoding and phylogenetic assignment of Haliclona *spp*

Among the samples collected, three morphotypes of *Haliclona* spp. Were identified with spicules concordant with, *H. cinerea*, *H. mediterranea,* and *H. oculata*, although the paucity of characters in *Haliclona* made our identifications very tentative. The first two species are common in the Mar Menor (Pérez-Ruzafa, 1989) and the third could correspond to *Haliclona* sp. recorded in the same work. To help identifying these *Haliclona* spp., the *COI* metabarcoding sequences from these samples were included and all other haplosclerids recovered in the metabarcoding analysis in a phylogenetic framework for the genus *Haliclona,* along with sequences of other species of *Haliclona* and other species from the order Haplosclerida from NCBI database (Supplementary Table 1). Clean sequences of *COI* from these samples through regular PCR could not be amplified, given the remarkable filtering activity of these sponges and retaining ability of foreign DNA, which produced clean bands but contaminated with many other organisms. Therefore, a fragment of the 18S ribosomal RNA (*18S*) was targeted for five representative specimens from the three haplotypes found (Ri2.2P, Ri2.3V, Ri1.3V, Urr2.2P and Urr1.2V), since it is a gene widely used in phylogenies of the order Haplosclerida (Redmond et al., 2007; Itskovich et al., 2007), and allows the comparison of these specimens with other *Haliclona* species from GenBank for which the *COI* is not available including the only *H. mediterranea* sequenced to date (GenBank ID: AY348879).

The eDNA extraction used for the metabarcoding was used to amplify the *18S* of the five representative specimens of the three different haplotypes (Ri2.2P, Ri2.3V, Ri1.3V, Urr2.2P and Urr1.2V). PCR reactions targeted a 1,045-bp fragment of the*18S* gene using the primers 1F (5’-GAAGAACCACCGTTGTTATTCAA-3’) and 5R (5’-ACC TCCR ATC T YCGG AT TAC A-3 ’) (Whiting, 2002) and consisted of a final volume of 13.5 µl containing 10 µl of 2× RedTaq PCR Master Mix (Thermo Scientific), 1 µM of each primer, 0.5 µl of BSA and 1 µl of template DNA (final concentration 80–200 ng µl^-1^). Due to their filtering habits and retention of foreign DNA, the extractions were diluted at a ratio of 1:500 to reduce the presence of foreign DNA and increase the probability of sponge amplification. The PCR conditions included an initial denaturation step at 95°C for 5 min was followed by 9 subsequent cycles of denaturation at 95°C for 30 s, annealing at 58°C, then decreasing 1°C per cycle for 30 s and extension at 72°C for 45 s, and then 30 cycles of denaturation at 95°C for 30 s, annealing at 50°C for 30 s and extension at 72°C for 45 s, followed by a final extension step at 72°C for 7 min. PCR products were purified with ExoSAP-IT (USB) and then sequenced on a 3730xl DNA Analyzer by Macrogen Inc.

Sequences of *COI* and *18S* were aligned separately using MAFFT (Katoh & Standley, 2013) as implemented in Geneious 10.1.3 (https://www.geneious.com; Kearse et al., 2012) with the default parameters, and then manually curated. The *COI* alignment was trimmed to 270 nucleotides and contained 47 sequences including the sequences of the present study along with published sequences of *Haliclona* species and other closely related taxa (Supplementary Table 1 and Figure 2). The *18S* alignment, spanning 1,045 nucleotides, was trimmed using Gblocks (Castresana, 2000) to select conserved blocks and then 935 nucleotides were selected for further analysis. The final alignment of the *18S* contained 43 terminals, including five newly generated sequences obtained in our study, as well as sequences belonging to published *Haliclona* species and other closely related taxa (see Supplementary Table 1 and Figure 1). All phylogenetic reconstructions were performed through the Maximum Likelihood (ML) method in RAxML implemented in raxmlGUI v.1.3 (Stamatakis, 2014), after selecting for the most likely nucleotide substitution model using jModelTest (Posada, 2008), and setting the search strategy to ML + through bootstrapping + consensus.

**Figure 2.**
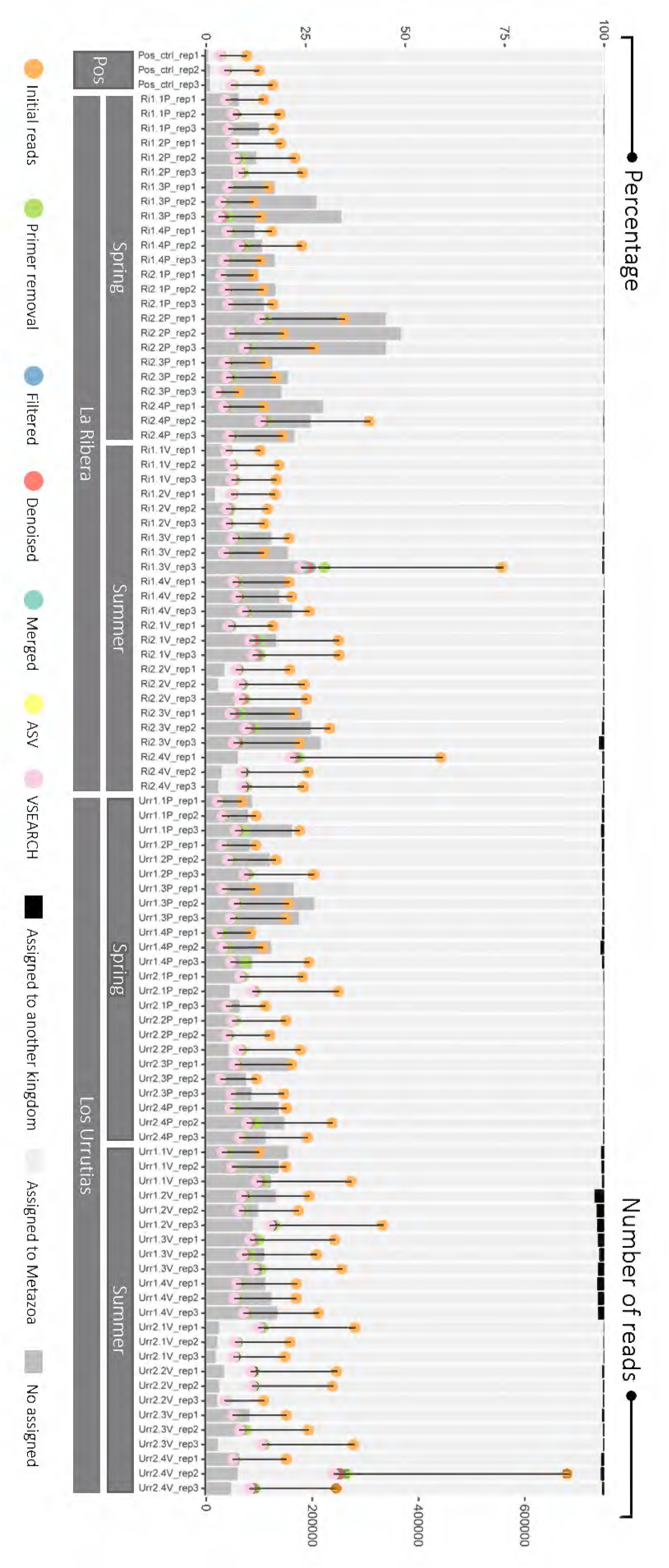
Combined graph showing both; the number of the kept reads for the different cleaning steps represented as circles of different colours, indicating the different steps of the bioinformatics pipeline (referenced to the marks of right axis and the colour code is shown in the legend) and the percentage of the cleaned reads after cleaning bioinformatic steps for each sample associated to three different levels: cleaned reads not assigned to neither kingdom (dark grey), cleaned reads associated to Metazoa kingdom (light grey) and cleaned reads associated to a kingdom but not metazoan (black) reads through the different cleaning steps. The different colours indicate the different steps of the bioinformatics pipeline and the code is shown in the legend, and the final kept reads after the cleaning steps are shown in pink (VSEARCH) along their taxonomic assignment strategy.

### Phylogenetic assignment of Porifera sequences

Some of the cleaned sequences that were taxonomically assigned to Porifera in the metabarcoding results were automatically assigned to sponge species from very distant locations, with low probabilities of being correct. Phylogenetic reconstructions are good practices to avoid misassigned sequences, sequences assigned to a low taxonomic level, or alignment with sequences not belonging to *COI* (Bidartondo, 2008; Meiklejohn, Damaso & Robertson, 2019). Therefore, phylogenetic analyses were carried out for sequences assigned to five different orders separately (Clionaida, Haplosclerida, Poecilosclerida, Spongillida and Suberitida) including specimens from GenBank and newly generated sequences in our lab (see Supplementary Table 1 and Figure 2). All alignments for each family of the class Demospongiae were carefully curated and then all phylogenetic reconstructions were performed as above.

### Statistical analyses

All diversity parameters were calculated for each sample by pooling technical PCR replicates and combining the values for the different taxa. The relative abundance of each taxon was used to calculate all diversity parameters and a presence-absence matrix was also used to calculate jaccard parameter of beta diversity. All estimations of diversity parameters were performed in *phyloseq* R package (McMurdie & Holmes, 2013). Alpha-diversity was estimated using Shannon’s diversity index with the *estimate_richness* function (Shannon, 1948) for each locality in each season. A two-way ANOVA was carried out with the locality (La Ribera and Los Urrutias) and season (spring and summer) as factors and the Shannon’s diversity index as the dependent variable calculate for each individual sampled. Pairwise mean differences were analysed using Tukey’s test with the *TukeyHSD* function of the *stats* package. Prior to ANOVA analyses, normal distribution and homogeneity of variances of the data were evaluated using *shapiro.test* and *bartlett.test* from *stats* package. Beta diversity was assessed by computing the Bray-Curtis distance matrix and Jaccard distance matrix (for binary data) using the *distance* function from the *phyloseq* package. Community differentiation between locality (La Ribera and Los Urrutias), seasons (summer and spring), their interaction and pairwise comparisons were examined through permutational multivariate analysis of variance (PERMANOVA) using the *adonis2* function from the *vegan* package, with both the Bray-Curtis and Jaccard distance matrix as the dependent variable. When significance differences were detected between groups, the multivariate homogeneity of group dispersions was tested with the *betadisper* function in the vegan R package to test that *adonis2* results are not due to differences in group dispersions. The community differentiation was visualized using both Bray-Curtis and Jaccard distance through principal coordinate analysis (PCoA), implemented with the *cmdscale* function in the *vegan* R package (Oksanen et al. 2013). To assess the completeness of sampling efforts, rarefaction and extrapolation curves were calculated using the *iNEXT* R Package (Hsieh et al., 2016), with bootstrapping set to 100.

## Results

### Phylogenetic assignment of host identity

For the host sequences collected as *Haliclona* spp., three different haplotypes were found as the dominant sequences (and hence considered as the host species) for the 313-bp fragment amplified for the *COI*: Hash_45e4b in all samples from La Ribera except for Ri22P, Hash_361bc in Ri2.2P, and Hash_7a781 in all samples from Los Urrutias (Supplementary Table 1). In the phylogenetic trees (Supplementary Figure 1 and 2), consistent results were found for the taxonomic assignment of the sequences, finding that they were not monophyletic for either *18S* or *COI* (Supplementary Figures 1 and 2), but clustered with the same species consistently in both trees. In the *18S* tree, the individuals Ri2.3V and Ri1.3V (Hash_45e4b haplotype) formed a well-supported clade together with the only sequence of *H. mediterranea* available in GenBank, and this clade was sister to *Chalinula molitba* and then *Callyspongia plicifera* (Supplementary Figure 1). In the *COI* tree (Supplementary Figure 2), the Hash_45e4b haplotype clustered with *C. plicifera*, but not *H. mediterranea* since there is no sequence available in GenBank for it. In this case, both Ri2.3V and Ri1.3V were considered to be *H. mediterranea* as well as the rest of individuals in the Ribera site, except for Ri2.2P, based on the dominant metabarcoding sequence in these samples and the concordance with the spicules showed by *H. mediterranea*. The individuals Urr2.2P and Urr1.2V and the Hash_7a781 haplotype, formed a well-supported group for both *18S* and *COI* with sequences belonging to the *H. cinerea* (Supplementary Figure 1 and 2), and therefore individuals with this haplotype were from now on considered as *H. cinerea*, as well as the rest of individuals in Los Urrutias, based on the dominant metabarcoding sequence in these samples and the concordance with the spicules showed by *H. cinerea*. Finally, the Hash_361bc haplotype (individual Ri2.2P) haplotype clustered with a clade composed of *H. urceolus* and *H. oculata* with strong support for *COI* tree (Supplementary Figure 2) and in the *18S*, in a robustly supported clade containing *Haliclona oculata* (Supplementary Figure 1). Therefore, given the similarity of both spicules and sequences with *H. oculata*, individuals with this haplotype (Hash_361bc) were henceforth referred to as *H. oculata*.

### Sequence read abundance and ASV/OTU richness

A total of 27,188,520 raw reads were obtained from the sequencing of the library with an average of 172,946.3±90,492.5 reads per sample. After cleaning, a total of 32.8±2.7% of reads were retained per sample. From these clean reads, 86.1±9.7% were assigned to a metazoan taxonomic category, 0.4±0.5% were assigned to another kingdom, and 13.5±9.6% were not assigned to any taxon from the GenBank and BOLD databases (Figure 2, Supplementary Table 1).

The distribution of the number of reads per species at each of the different sample sites was similar across samples, with a predominantly low number of reads for the majority of the species (around 40–50% of the total number of species) and a few species with very high values (Supplementary Figure 3). This pattern appeared to be somewhat attenuated at locality 1 of Los Urrutias in the spring, where the distribution was less skewed(Supplementary Figure 3).

### Phylogenetic assignment

Given the importance of the phylum Porifera in terms of taxa detected and relative abundance in the study, as well as the probability of detection of invasive sponge species, an in-depth analysis of some of the sequences with dubious assignments within this phylum was performed (i.e., those with assignments to either family or order). Although the BLAST results; resulted in four different sequences assigned to *Poecilosclerida* sp., these sequences were grouped with *Mycale fibrexilis* in the phylogenetic analysis and therefore considered as *Mycale fibrexilis* (*b*=92; Supplementary Figure 6). Within the order Spongillida, a sequence was found that was initially assigned to the genus *Ephydatia* Lamouroux, 1816, a genus found only in freshwater ecosystems. The phylogenetic analysis, grouped this sequence with different spongilid species including *Metschnikowia tuberculata* Grimm, 1877 (Supplementary Figure 7), the only known species of this order inhabiting marine and brackish ecosystems (Sokolova et al., 2020). However, given the uncertainty in the species assignation for this sequence, we simply named this sequence as Spongillida sp. (Supplementary Figure 7). Within the order Haplosclerida, besides the host sequences, sequences that clustered with *Haliclona xena*, *Haliclona elegans*, *Haliclona* aff. *aquaeductus*, *Haliclona cinerea* were retrieved, and an unidentified species of *Haliclona* (Supplementary Figure 2).

Within the order Suberitida, we obtained two different sequences that were only assigned to Demospongiae. One of these sequences clustered robustly with several sequences of *Halichondria bowerbanki*, which is the reason why we assigned this sequence to this species (Supplementary Figure 8). The other sequence clustered with unidentified species of *Pseudosuberites* and other halichondriids, and we named it as Halichondriidae sp. (Supplementary Figure 8). In addition, we retrieved a sequence that formed a well-supported clade with three *Suberites* spp.: *Suberites* sp., *S. aurantiacus* and *S. diversicolor*, and therefore that sequence was named as *Suberites* sp. (Supplementary Figure 8).

Finally, a phylogenetic tree was also built for two sequence blasting with the family Clionaidae. One sequence appeared in a well-supported clade with different *Pione* species including the putatively invasive species *Pione vastifica* and *P. lampa*; this sequence was thus named *Pione* sp. (Supplementary Figure 9). The other sequence did not cluster with any sequenced species, and therefore we named as Clionaidae sp. (Supplementary Figure 9).

### Alpha and beta diversity

In spring, the site Los Urrutias (3) showed the highest alpha diversity value and the lowest variance within the different sample sites (Table 2; Figure 3A). The results of the Shapiro-Wilk test (*W* = 0.96, *p* = 0.3) and Bartlett’s test (*Bartlett’s K-squared* = 10.51, *df* = 7, *p* = 0.16) were not significant, indicating a normal distribution and homogeneity of variance, respectively, and the suitability of conducting parametric analyses. The ANOVA performed on the alpha diversity index revealed slightly significant differences between localities (La Ribera *vs* Los Urrutias), but not between seasons (spring *vs* summer), or the interaction between region and season (Table 2).

**Table 2.** Two-way ANOVA conducted with the Shannon index as the dependent variable, considering both Locality (La Ribera and Los Urrutias) and Season (spring and summer) as factors. The table presents the Degrees of Freedom (df), Sum of Squares (SS), Mean Square (MS), F-statistic (F), and P-value (p) associated with the analysis. Statistically significant results p < 0.05 in bold.

|  | Df | SS | MS | F | *p* |
| --- | --- | --- | --- | --- | --- |
| Locality (La Ribera and Los Urrutias) | **1** | **1.15** | **1.15** | **4.37** | **0.045\*** |
| Season (spring and summer) | 1 | 0.34 | 0.33 | 1.29 | 0.265 |
| Locality*Season | 1 | 0.83 | 0.83 | 3.17 | 0.085 |
| Residuals | 28 | 7.37 | 0.26 |  |  |

**Figure 3.**
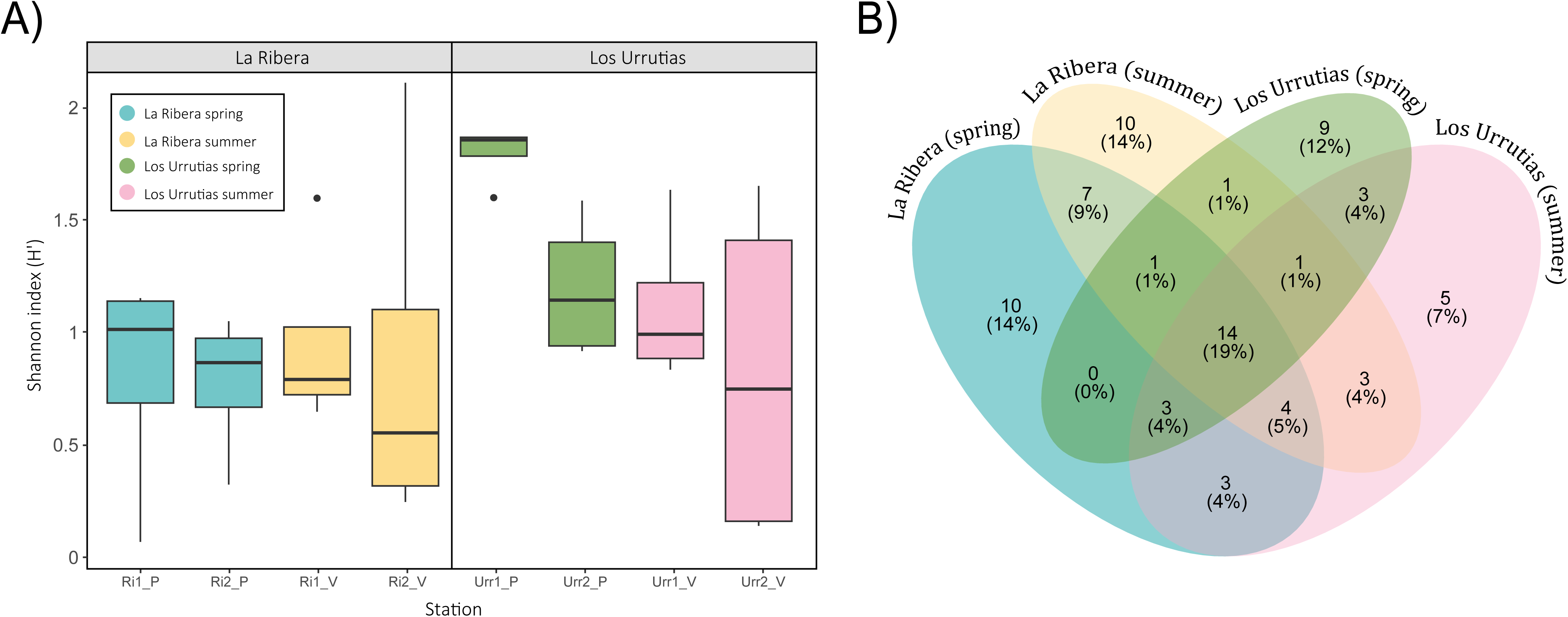
A) Box plot of alpha diversity for species-level richness based on Shannon alpha diversity index for eDNA samples for the four sites sampled in both seasons. B) Venn diagram of the shared species for the two locations (La Ribera and Los Urrutias) in the two seasons (spring and summer).

For beta-diversity, the results of the Principal Coordinates Analysis for the Bray-Curtis indices considering all samples, clearly separated samples from La Ribera and Los Urrutias, with PCoA axes 1 and 2 explaining 38.9% and 15% of the variance, respectively for Bray-Curtis and 19.2% and 13% for Jaccard, reppctively (Figure 4.A, Supplementary Table 2). Los resultados tanto para la matriz de distancias de Bray Curtis y de Jaccard fueorn congruentes en usmayoria para los estadicticos de PERMANOVA y betadisp. The differentiation between La Ribera and Los Urrutias was also supported by the PERMANOVA results of both matrixes, with significant differences between the regions and both slight and significant differences in the interaction between region and season for BrayCurtis and Jaccard respectively (Figure 4.B, Supplementary Table 3). Para la matriz deJaccard tabn encontramos diferencias significativas entre estaciones. In addition, the PERMDISP test rejected any significant differences in the dispersions between groups in any of the three cases for both matrixes (region, season and region:season, Figure 4B, Supplementary Table 2).

**Figure 4.**
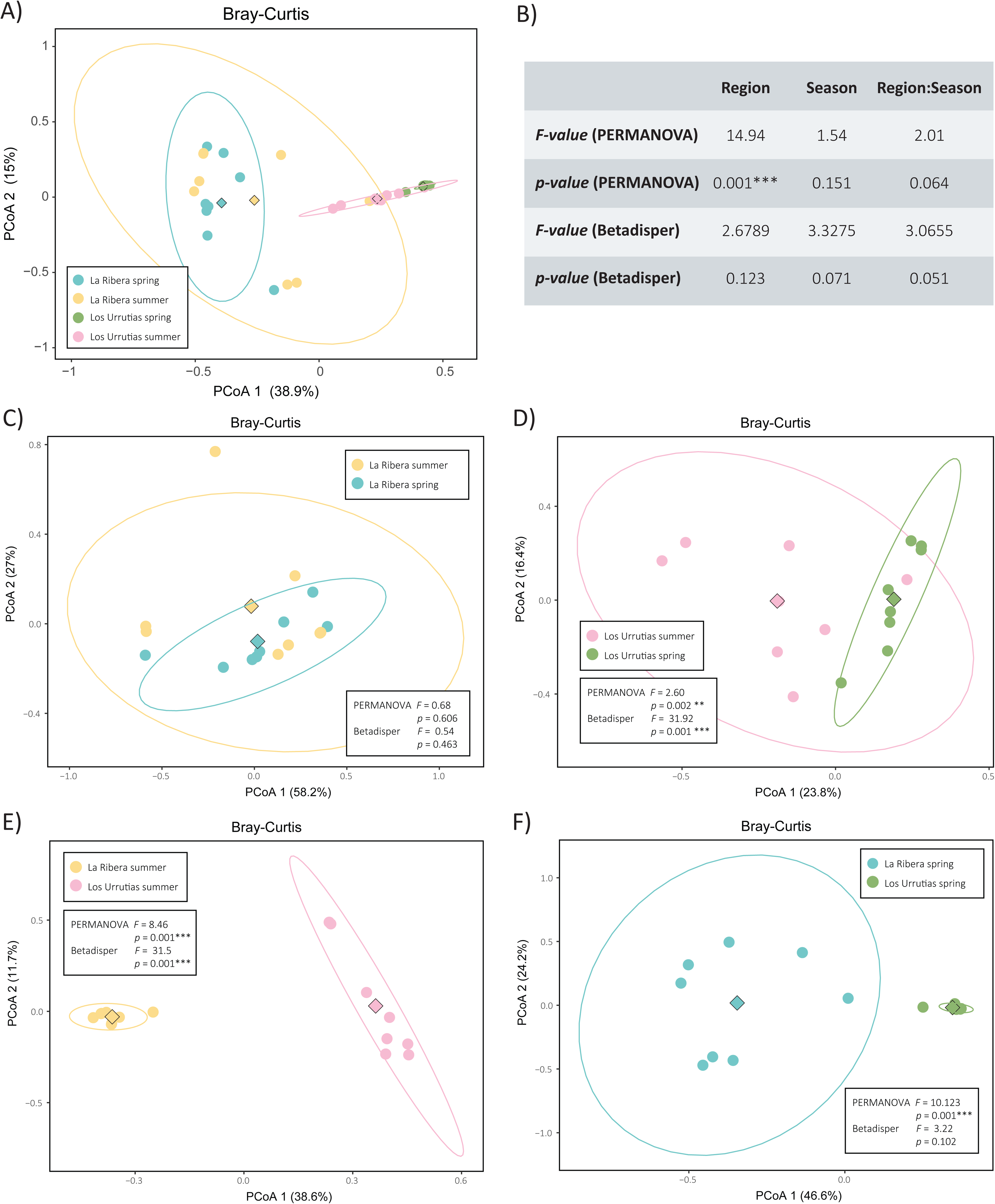
Principal Coordinates Analysis for Bray-Curtis indices and summary of the results obtained for both PERMANOVA and Betadisper analyses. The centroids are marked as a diamond. A) PCoA of all samples (see details in legends). B) Summary table of PERMANOVA and Betadisper for all samples. The following figures include PCoA (see legend for details), including summary of PERMANOVA and Betadisper results for C) La Ribera in summer and spring, D) Los Urrutias in summer and spring, E) La Ribera and Los Urrutias in summer, F) La Ribera and Los Urrutias in spring.

In pairwise comparisons within the same locality, no significant differences were found for samples from La Ribera in spring and summer (Figure 4C, Supplementary Table 2) but differences were detected in Los Urrutias for Bray-Curtis and Jaccard (Figure 4D, Supplementary Table 2). In the last case, although no homogeneity of variance was detected only for Bray-Curtis matrix, there were obvious differences in community composition between seasons, especially along the first axis, which explained 23.8% of the variance (Figure 4D, Supplementary Table 2). When comparing sites within the same season, significant differences were detected between La Ribera and Los Urrutias in both summer (Figure 4E, Supplementary Table 2) and spring (Figure 4F, Supplementary Table 2).

These differences in community assemblages included remarkable differences in the relative abundances of specific taxa. In spring, relative abundance was more or less evenly dominated by either Annelida or Porifera reads in all replicates from La Ribera, while in Los Urrutias, Annelida was the only phylum that dominated the majority of reads across all replicates: *Haplosyllis* sp., *Branchiomma boholense* and *Terebella lapidaria* (Figure 5A and 6). In the summer, these relative abundances shifted further, with La Ribera then dominated by Annelida reads in most replicates, while Los Urrutias showed greater heterogeneity in the relative read abundance compositions (Figure 5A and 6). In both areas, but especially in Los Urrutias in summer, several replicates were dominated by the cnidarian *Kirchenpaueria* sp., and in Los Urrutias, one replicate detected an increase in Arthropoda-associated reads (Figure 5A and 6). At the species level, further noticeable differences included high abundances of *Haplosyllis* sp. and low abundances of *Branchiomma boholense* in La Ribera, as well as the presence in both seasons of *Branchiomma bairdi* in La Ribera, while in Los Urrutias we only detected high abundances of *Branchiomma boholense* (Figure 6).

**Figure 5.**
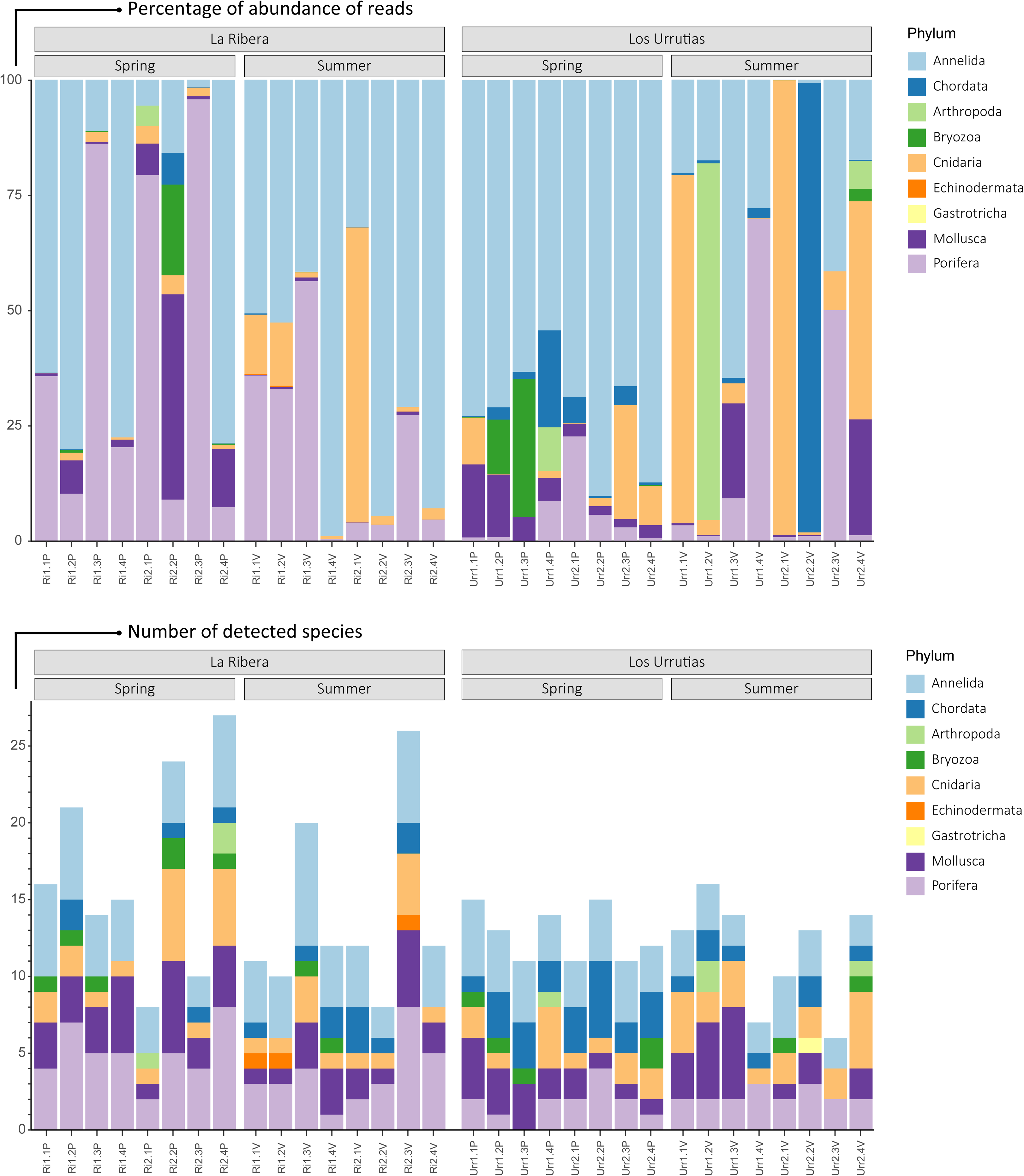
Barplot showing the number of taxa detected at each phyla from each biological sample (PCR replicates are combined) and their relative abundance. A) Percentage of reads obtained for each phylum for each sample. B) Number of species detected for each phylum in each sample.

**Figure 6.**
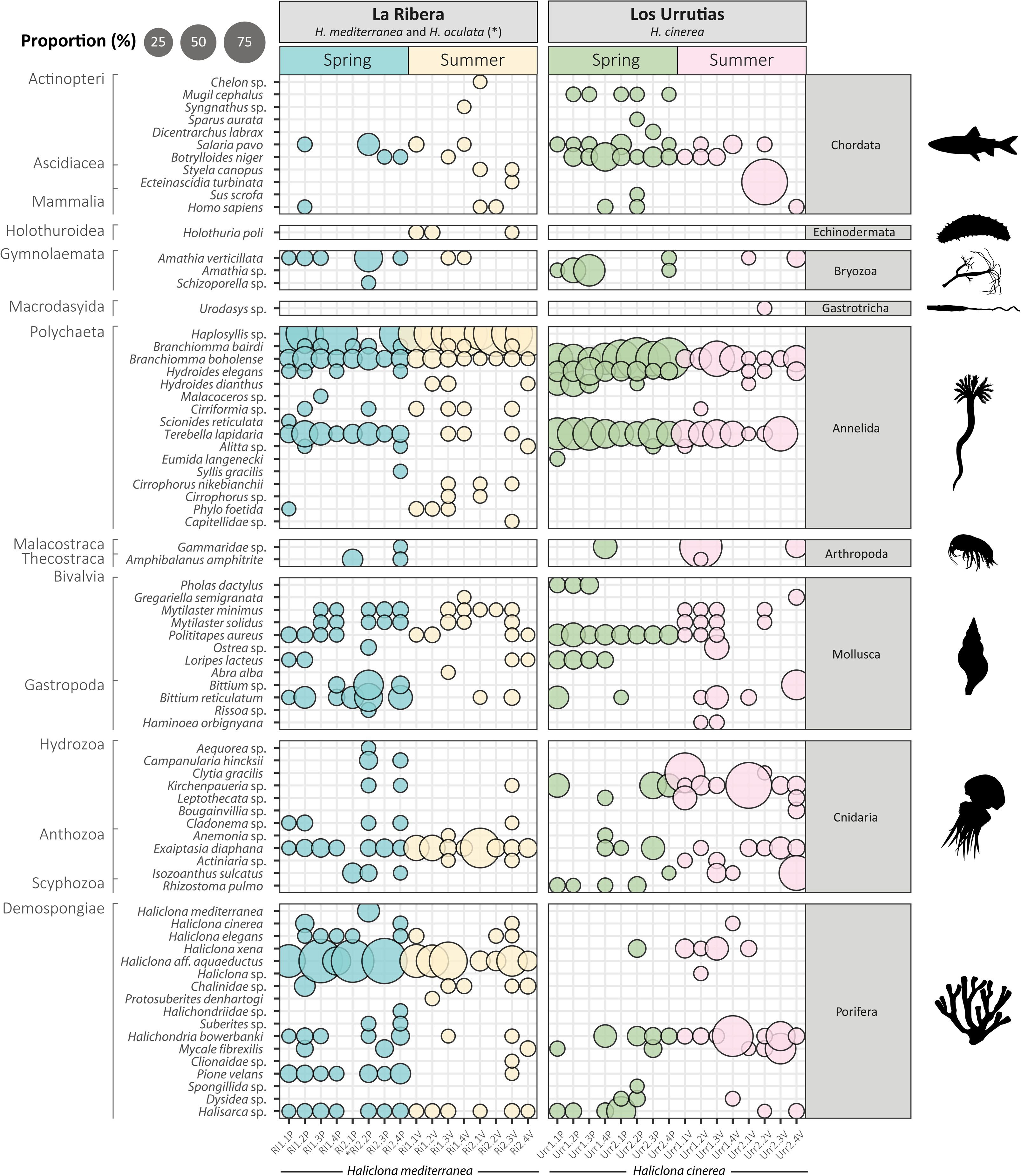
Bubble plot of taxa detected using sponge nsDNA in both regions and seasons for each phylum. Circle size indicates the proportion of reads assigned to each taxon per sample. Colours represent the type of samples: blue for La Ribera in spring, yellow for La Ribera in summer, green for Los Urrutias in spring and pink for Los Urrutias in summer.

### Biodiversity composition and structure

A total of 76 taxa were detected at species, genus, family and order level belonging to the phyla Porifera (18), Annelida (16), Cnidaria (12), Mollusca (12), Chordata (11), Bryozoa (3), Arthropoda (2), Echinodermata (1), and Gastrotricha (1) (Figure 6). In most samples, Annelida was the group with the highest relative abundance of values obtained (average of 52.16%), followed by Porifera (22.10%), Cnidaria (12.84%) and Mollusca (7.08%) (Figure 5 and 6; Supplementary Table 2). Only one species of Echinodermata appeared in the three biological samples in La Ribera during the summer season, with a relative abundance of <1%. The phylum Gastrotricha appeared in only one biological sample from Los Urrutias during the summer season, with a relative abundance percentage of <1%. Although very low numbers of species were detected for crustaceans, high relative abundances of reads were detected at Los Urrutias in summer (Urr1.2V), as well as for Cnidaria and Chordata phyla at Los Urrutias in summer (Urr1.1V, Urr2.1V, Urr2.2V and Urr2.4V). There was an average of 3.81 species detected per sample for the phylum Annelida, followed by 3.15 species for the phylum Porifera, 2.06 species for the phylum Cnidaria and 1.4 species for the phylum Chordata.

Fourteen taxa were consistently detected at both localities in both seasons (Figure 3B, Supplementary Table 2) and included: *Halichondria bowerbanki*, *Halisarca* sp. and *Mycale fibrexilis* (Porifera); *Exaiptasia diaphana* and *Kirchenpaueria* sp. (Cnidaria); *Bittium reticulatum* and *Polititapes aureus* (Mollusca); *Alitta* sp., *Terebella lapidaria* and *Branchiomma boholense* (Annelida), *Amathia verticillata* (Bryozoa) and *Salaria pavo*. (Chordata); *Botrylloides* sp. and *Homo sapiens* (Chordata). More specifically, four taxa were found in all biological samples for both seasons: *Halichondria* sp. (Porifera), *Exaiptasia diaphana* (Cnidaria) and *Branchiomma boholense* and *Terebella lapidaria* (Annelida). Within the same locality for the different seasons, a total of 26 taxa (46% of the total) and 21 taxa (45% of the total) were shared in La Ribera and Los Urrutias, respectively (Figure 3.B). Only between 5 and 13% of the detected species were uniquely found in one site and one season (Figure 3). Some of the species detected consistently across samples in La Ribera in summer but not anywhere else were the annelid *Cirrophorus nikebanchii* and the echinoderm *Holothuria poli* (Figure 5 and 6). In Los Urrutias, during the spring, we detected the gastropod *Pholas dactylus* and the jellyfish *Rhizostoma pulmo* (Figure 6).

### Rarefaction and consistency in taxa detection

Rarefaction curves showed that sequencing depth was sufficient to fully recover biodiversity present in all PCR replicates of eDNA (Supplementary Figure 4), while curves of species richness represented by replicates in the respective sampling areas suggest that some richness was not recovered by the number of replicates employed here (Supplementary Figure 5).

The efficiency and consistency of our methodology for taxa detection across our PCR and biological replicates was explored (Supplementary Figure 6). First, it was tested whether the same taxa were being recovered by the different PCR replicates (three technical replicates for each eDNA tube extracted), and it was found that in 46% of the cases all the recovered taxa were detected by all three PCR replicates, in 17% of the cases by two PCR replicates and by only one PCR replicate in 38% of the cases. Then, it was tested whether the same taxa could be recovered from the different biological samples collected at each sampling site (four biological samples per site) and separated by season, and found that only in 14% of the cases all the recovered taxa were detected by the four biological samples, in 17% of the cases the species was detected by three biological samples, in 21% of the cases the species was detected by two biological samples and in 48% of the cases the species was detected by one biological sample. Then, it was assessed whether the results were consistent across sites within the same area during the same season, and in 56% of the cases, the taxa was detected at only one of the two sampling points, while in 45% of the cases, the taxa was detected at both sampling points.

## Discussion

The Mar Menor is a highly heterogeneous coastal lagoon characterised by complex and diverse faunal communities, with few dominant species but high overall abundances (Pérez-Ruzafa et al. 2005, 2011, 2019a, 2020). Approximately, up to 40% of the species present can vary annually (Pérez-Ruzafa et al. 2019a), making biodiversity monitoring both challenging and costly due to significant spatial and temporal variation. In addition, these ecosystems are heavily influenced by human activities and climate change, leading to significant shifts in environmental conditions, including changes in salinity, temperature (Vergara-Chen et al. 2010; González-Wangüemert & Pérez-Ruzafa 2012), and nutrient inputs (Pérez-Ruzafa et al. 2005), ultimately affecting the viability, reproduction and survival of many species.

Our approach using natural samplers to collect eDNA provided valuable insights into biodiversity of this area, enabling the identification of common and abundant species associated to small piers of the lagoon, as well as of some important allochthonous and commercially important species (Verdiell-Cubedo et al., 2013; Giménez-Casalduero et al., 2016; Terradas-Fernández et al., 2022). Importantly, we also found some seasonal variability in community composition, and consistent spatial differences across seasons, consistent with the changing environmental conditions known to occur in this lagoon (Pérez-Ruzafa et al., 2007; Roman et al., 2009).

### Biodiversity composition and structure

The severe degradation that the Mar Menor has experienced as a result of strong anthropological pressures has affected both the composition and the density of the biodiversity in this lagoon (Pérez-Ruzafa et al., 2005, 2006; Sala-Mirete et al., 2023). Monitoring the dynamics of biodiversity composition and structure has then become urgent for conservation purposes. The opportunistic subsampling of individual sponges for eDNA analysis enabled us to produce a general view of the composition of local biodiversity, providing key information and an efficient method for assessing the biodiversity in a system as vulnerable and dynamic as the Mar Menor. Here, 74 metazoan taxa belonging to 9 different phyla were detected using the metabarcoding approach, excluding the common vertebrates *Homo sapiens* and *Sus scrofa*, which in the case of humans originates from various sources, including operational contamination of samples, waste run-off, and/or the bathing public. Notably, the nsDNA technique used in this study identified significantly more species than a previous traditional taxonomic survey based on morphological characterisation, which identified 31 species in 7 different phyla (Sandoninni et al., 2021). Among these species, 15 taxa detected by sponge nsDNA had not been previously reported from the Mar Menor, representing approximately 21% of the total list of species recorded in the lagoon. These newly detected taxa include: the ascidian *Botrylloides niger* Herdman, 1886, the gastrotichean *Urodasys* Remane, 1926, the annelids *Cirrophorus nikebianchii* Langeneck, Barbieri, Maltagliati & Castelli, 2017, *Eumida langenecki* Teixeira, Vieira, Ravara, Costa & Nygren, 2022 and *Branchiomma bairdi* (McIntosh, 1885), the molluscs *Mytilaster solidus* Monterosato, 1883 and *Gregariella semigranata* (Reeve, 1858), the cnidarians *Cladonema* sp. Dujardin, 1843, *Campanularia hincksii* Alder, 1856, *Aequorea* sp. Péron & Lesueur, 1810 and the sponges *Halisarca* sp. Johnston, 1842, Spongillida Manconi & Pronzato, 2002, *Pione velans* (Hentschel, 1909), *Protosuberites denhartogi* van Soest & de Kluijver, 2003 and *Haliclona cinerea* (Grant, 1826). Another new detected species is *Isozoanthus sulcatus* (Gosse, 1860), at present in taxonomic controversy with the possibility to belong to the family Nanozoanthidae (Fujii & Reimer, 2013).

Other taxa not previously recorded in the Mar Menor were; the annelids *Alitta* sp. Kinberg, 1865, *Terebella lapidaria* Linnaeus, 1767, Scionides reticulta (Ehlers, 1887) and *Haplosyllis* sp. Langerhans, 1879 and the sponges *Mycale fibrexilis* (Wilson, 1894), *Halichondria bowerbanki* Burton, 1930, *Haliclona aquaeductus* (Schmidt, 1862) and *Haliclona xena* de Weerdt, 1986. However, the taxonomic assignment of these taxa remains uncertain due to the lack of genetic data for closely related species reported in the region. This limitation in genetic resolution prevents more accurate identification and highlights the need to expand DNA sequence repositories.

Given that sponges play a fundamental role in serving as host organisms for a wide variety of marine animals (Wendt et al., 1985; López et al., 2001), it can be expected that a greater amount of biomass, and therefore eDNA, of species that inhabit sponges will be filtered by the sponges. Thus, it is common for the detection of eDNA to be biased by ecological factors, resulting in much of the eDNA detected by sponges originating from species that inhabit them (Jeunen et al., 2023). In this sense, the data collected here are consistent with previous studies that have analysed microfauna and found annelid species to be the most frequent and abundant taxa living in and around sponges (Wendt et al., 1985; Martin & Britayev, 1998; Lopez et al., 2001; Krauter et al. al., 2006; Jeunen et al., 2023), as well as with other surveys on eukaryotic biodiversity assessment through eDNA collected from sponges (Gallego et al., 2024; Jeunen et al., 2023). However, the results are congruent with studies on trophic webs (Pérez-Ruzafa et al., 2020) and other recent studies on the assessment of the biodiversity of the subcoastal bottoms of the Mar Menor, carried out through the survey of artificial settlement structures (Sandonnini et al., 2021). Depending on whether the estimations are made on biomass or number of individuals and the type of community sampled in general Annelida, Mollusca, Arthropod and Cnidaria are the most abundant and diverse phyla detected by traditional surveys (Pérez-Ruzafa et al., 2020, Sandonnini et al., 2021), Annelida being the most diverse (Pérez-Ruzafa and Marcos, 1993; Sandonnini et al., 2021), which is also the main result of this study, with the phylum Annelida being the most represented, with up to 60% of the total reads, 10 species and 6 additional OTUs identified at genus level (Figure 5).

Besides annelids, we found a noticeable abundance of sponges, cnidarians and molluscs but very few echinoderms and arthropods. Despite sampling hard substrate community, this low diversity of arthropods was surprising, since previous surveys based on traditional methods had indicated that this phylum was the one of the most abundant in the Mar Menor lagoon, mainly the orders of Amphipoda and Isopoda (Pérez-Ruzafa, 1989; Pérez-Ruzafa et al., 2020; Sandoninni et al., 2021). Differences in the relative presence and/or abundance of different taxonomic groups between methods can be explained by the morphology and anatomy of the taxa, in addition to biased PCR approximations (Kelly et al., 2019). It has previously been hypothesised that the presence of an exoskeleton, characteristic of arthopods, reduces the release of DNA into the environment, which may affect its detection in eDNA-based studies (see Shokralla et al., 2010; Zizka et al., 2018; Kestel et al., 2024). Therefore, we consider that, although there is a possibility of a reduction in the representation of the phylum Arthropoda in the Mar Menor, this fact may be more influenced by the low DNA density for this phylum in the environment as a consequence of the presence of exoskeleton. Interestingly, we also provide the first record of the phylum Gastrotricha in the Mar Menor, which though a novel finding, is unsurprising given that there has been little focus on the local meiofauna beyond some previous research into the lagoon’s foraminifera, nematodes and amphipods. The reason why other meiofaunal dwellers such as for instance Nematoda, one of the most abundant organisms in this habitat (e.g. Barnes et al., 2008), were not also detected remains unknown.

As for echinoderms, their species richness has historically been considered to be low in the Mar Menor (Pérez-Ruzafa et al. 2005), likely due to their limited adaptive capabilities to high salinity conditions (Lawrence 1990; Vergara-Chen et al. 2010). In the 1980s, the only common species was the ophiuroid *Amphipholis squamata* (Delle Chiaje, 1828) and in a scarce density and more limited distribution the species *Oestergrenia thomsonii* (Herapath, 1865), *Paracentrotus lividus* (Lamarck, 1816), *Holothuria impatiens* and *Oestergrenia digitata* (Pérez-Ruzafa, 1989; González-Wangüemert, et al., 2018). Since the 1980s, *Holothuria poli* (Delle Chiaje 1823) colonized and quickly spread in the lagoon, which has been shown to be one of the most adaptable species of echinoderms, and currently being the most abundant echinoderm in the lagoon (Pérez-Ruzafa and Marcos, 1993; Vergara-Chen et al., 2010). It is noteworthy that in our study we only detected eDNA of *H. poli* in La Ribera during the summer. This period coincides with its spawning season of *H. poli*, when large numbers of sperm are released into the environment (Navarro et al., 2012; Bardanis & Batjakas, 2018). Therefore, given its sedimentary habits, it is likely that the detectability of this species is highest in summer, as a consequence of the high load of genetic material released into the water column during reproduction. BUSCAR INFO DE REPRODUCCION DE ANELIDO

We detected significant differences in the structure of the biodiversity across spatial and temporal scales, except for the site La Ribera, which remained more stable across seasons. In the Mar Menor, environmental parameters such as salinity and nutrients change seasonally (even at fortnight scale) and spatially, and biological factors, such as chlorophyll α concentration change seasonally, which produces changes in biodiversity composition and abundance in this and other Mediterrenean areas including others lagoons (Pérez-Ruzafa et al. 2007; Balata et al. 2006; Lucena-Moya et al. 2012; Khedri et al. 2017).

### Species of conservation concern

The monitoring of biodiversity is a fundamental tool that provides underlying information about the health of ecosystems and how they are responding to environmental changes and human activities (Cordier et al., 2021). Within the broader pattern of biodiversity assessed here, we were able to detect the presence and relative abundance of several species of conservation concern in the Mar Menor lagoon, which are considered of interest because of their vulnerability, importance in commercial fisheries or invasive nature.

#### Species of commercial interest

Among the fish species of commercial interest detected in this study, we found *Chelon* sp. (being *C. auratus, C. labrosus, C. ramada* and *C. saliens* the most abundant species of the genus *Chelon* in the lagoon), *Mugil cephalus* and *Spaurus aurata* which constitute the main species in the Mar Menor fisheries (Marcos et al. 2015). The detection of these species is congruent with previous studies where Mugilidae taxa and *Sparus aurata* are the most common and abundant taxa in the shallow areas of the Mar Menor, a trend that seems to be maintained since 2002 (Guerrero-Gómez et al., 2021). It should also be noted that Guerrero-Gómez et al. (2021) detected *Atherina boyeri* as a common and abundant species, which was not detected in the present study.

#### Threatened species

In our study, we found some species of conservation interest with different levels of threat. We detected reads associated with the genus *Syngnathus*, a genus reported in the lagoon with the species *S. thyple* and *S. abaster* the most frequently detected in the Mar Menor (Pérez-Ruzafa, 1989; Guerrero-Gómez et al., 2021). We also found reads assigned to the molluscan *Pholas dactylus*, an endangered species included in Appendix II of the Convention on the Conservation of European Wildlife and Natural Habitats (Bern Convention). Populations of this species in the Mar Menor are common in the compacted red clays habitats located in the south-western area of the lagoon (Pérez-Ruzafa, 1989; Pérez-Ruzafa et al., 2020). Recently it has been recorded that some populations have been decimated and were thought to be locally extinct following recent episodes of eutrophication and hypoxia, before a small population was recently discovered on the shallow white clay seabed (Giménez-Casalduero et al. 2016). Interestingly, other threatened but still abundant species in the Mar Menor are *Pomatoschistus marmoratus* (Risso, 1810) and *Apricaphanius iberus* (Valenciennes, 1846) (Guerrero-Gómez et al., 2021), but these were not detected through our eDNA technique although this could be due to the habitat preferences of these species, which are more frequent on muddy and seagrass bottoms.

#### Allochthonous and invasive species

Changes in the physico-chemical conditions of the Mar Menor, caused by various human activities and exacerbated by climate change and maritime traffic, have reduced the barriers that prevented the entry and establishment of many alien species to the Mediterranean, leading to an increase in these species in recent decades (Pérez-Ruzafa et al., 2008). The spread of ocean sprawl, in the form of pillars, piers, jetties and sea walls?, although potentially allowing the recovery of certain taxa by providing hard substrate to colonise, also allows the introduction and spread of invasive species (Glasby et al. 2007). In addition, the high dynamism of lagoon conditions, related to both anthropogenic and natural factors, and the heterogeneity of faunal communities in the Mar Menor make it difficult to establish clear categories of allochthonous species (León et al., 2016), and it is also a challenge to determine the impact that those allochthonous species may have on native species (Ruiz et al., 1997). Therefore, the development of long-term biodiversity monitoring strategies is one of the best tools we can use to increase information and knowledge about the composition of an ecosystem (Cordier et al., 2021). Here we detected some species previously reported as allochthonous or invasive and already established in the lagoon, such as the ascidian *Styela canopus* (Gonzalez-Carrión, 2015) and the arthropod *Amphibalanus amphitrite* (Pérez-Ruzafa, 1989), the latter considered as invasive in the Mediterranean (Molnar et al., 2008), and whose first citations in the Mar Menor were in 1989 and 2015 respectively. We also detected the jellyfish species *Rhizostoma pulmo*, which is not considered strictly allochthonous, as this species colonised the lagoon after the widening of el Estacio inlet and since the 1990s, has produced annual proliferations (Pérez-Ruzafa et al. 2002; Fernández-Alías et al., 2020). Notably, since 1990s, massive proliferations of this cnidarian have been reported, causing important socio-economic impacts (Prieto et al., 2010; Fuentes et al., 2011). Additionally, we detected the presence of the sponge *Haliclona oculata*, considered an allochthonous species that is occasionally found on the hulls of recreational boats (González-Carrión, 2015; Casalduero et al., 2016), and the ascidiacean *Botrylloides niger*, a highly invasive species reported from all oceans, which is believed to be native to the West Atlantic (Sheets et al., 2016), and only confirmed in Turkey within the Mediterranean (Temiz et al., 2023).

We also detected two invasive annelid species of the genus *Branchiomma* (*B. boholense* and *B. bairdi*) from the Polychaeta class, one of the animal groups with the highest number of established and invasive taxa in the Mediterranean compared to any other sea in the world (Çinar, 2013). Concretely, the genus *Branchiomma* is of particular concern, as they can reach very high densities from very few colonizing individuals (Tovar-Hernandez et al., 2011). In the Mar Menor lagoon the first record of the species *Branchiomma boholense* was in 2009 (Roman et al., 2009), which was also the first record in the Iberian Peninsula. In the present work we have detected the presence of *B. boholense* in both sampling stations as well as in both spring and summer. The high detectability of this species supports previous findings by Roman et al. (2009), who found dense populations of *B. boholense* in the Mar Menor. The difficulty in distinguishing *B. boholense* from *B. bairdi* based on morphological features has led many experts to question the accuracy of species identification (Evans et al., 2015). However, despite their great morphological similarity, there is a marked genetic differentiation between them (Del Pasqua et al., 2018). In the Mar Menor, the identification of these species has been controversial. Initially, these populations were described as *B. boholense* based on morphological features (Roman et al., 2009), later reclassified as *B. bairdi* (Arias et al., 2013; Giménez-Casalduero et al., 2016), and a subsequent publication using molecular data confirmed that the population belonged to the species *B. boholense* (Del Pasqua et al., 2018). Therefore, with the results obtained, we support the coexistence of both species in the Mar Menor lagoon. Similarly, *B. bairdi* was only detected in all samples from La Ribera locality, regardless of the season; the absence of reads associated with *B. bairdi* in Los Urrutias could indicate the absence of this species here or a smaller population than that of *B. boholense*, which is widely distributed throughout the lagoon.

### Applications, limitations and future directions

As eDNA analysis establishes itself as a mainstay of marine ecological studies, the opportunistic use of abundant, readily available, resilient, “natural samplers” may also play an increasingly significant role in certain contexts. As we have seen in this instance, abundant *Haliclona* spp. can be used to obtain informative data on marine biodiversity, with a focus on the community features around artificial structure (see also Cai et al. 2024). There were nonetheless limitations to the scope of this research. Biodiversity monitoring through eDNA can involve notable challenges, such as when assessing presence versus viability of populations, PCR artifacts and primer biases, or the viability of eDNA metabarcoding as a quantitative tool (Goldberg et al., 2015; Sigsgaard et al., 2020). Natural eDNA samplers pose their unique challenges, such as variance in the degradation and retention of eDNA within different types of sampler tissue, or PCR resource competition between eDNA and the DNA of the host (i.e., the natural sampler), which can result in low representation of species represented in eDNA (e.g., Weber et al. 2023). Previous studies have also shown that the capacity of sponges to retain eDNA within their tissue varies across species, and that this variation can result in differing characterizations of local biodiversity based on sponge eDNA (Brodnicke et al., 2023; Cai et al., 2023, Neave et al. 2023, Gallego et al. 2024). In the present study, this phenomenon could have influenced the variation of biodiversity composition between the two sites sampled by collecting several species of the genus *Haliclona* as hosts, rather than just one species. However, these *Haliclona* species are very similar in their filtration capacities and microbiome composition (all being Low Microbial Abundance sponges), and potentially they could retain similar amounts of eDNA. Nonetheless, in our study, although the use of nested replicates should ensure maximum taxa detection and minimum bias, we found variability between taxa detected for different samples at the same level, even when assessing the same host species. This is consistent with previous literature, which has shown that species present at low abundances exhibit greater stochasticity in detection through eDNA metabarcoding (Shaw et al., 2016) and reflects on the importance of replication. Additionally, we hypothesise that the high percentage of host reads may further increase this stochasticity, although further studies are required to confirm this.

Importantly, as it is often the case with eDNA studies, the quality of taxonomic identifications here is limited by some fundamental issues. These include the type of genetic marker used in metabarcoding, the existence of reference sequences in the database for the species sampled, and that the sequences loaded into the database are assigned to the correct species. The most commonly used genes for genetic monitoring of metazoans are the cytochrome oxidase I gene (*COI*) and the mitochondrial 18S rRNA gene (*18S*) (Leduc et al., 2019; Clarke et al., 2021; Di Capua et al., 2021; Zhao et al., 2021). To date, *COI* has certain advantages over *18S*, as *COI* is better at offering species-level resolution than *18S* (Giebner et al., 2020; Tang et al., 2012). This is mainly due to a better reference database for *COI* than for *18S*, and because *18S* has a slower rate of evolution than *COI*, which can sometimes make it difficult to distinguish between different species (Othman et al. 2021). In this study, we managed to identify 57.89% of the reads at the species level and 32.89% at the genus level, with only 9.21% of the specimens identified at higher taxonomic levels and a total of 14% of reads were unassigned at any taxonomic level. Some of the species detected in this study and designated as new records should be treated with caution. This is because, although the *COI* is the universal marker to metazoan DNA taxonomy (Hebert, Cywinska, Ball, & DeWaard, 2003; also see the International Barcode of Life Project, iBOL; http://www.ibol.org/), there are species that have not been sequenced for this gene, and although the bioinformatic pipeline adjusts taxonomic ranks based on the percentage of identity obtained for each taxonomic assignment, the lack of genetic information for certain species may lead to incorrect taxonomic classifications. Therefore, while this study proposes new species detected in the Mar Menor, these should be confirmed using additional sampling methods, since some species could have been assigned to a nearby species.

In future work, the combined use of multiple genetic markers would certainly aid in better biodiversity recovery (Stat et al., 2017; Compson et al., 2020; Gallego et al., 2024); nonetheless, a strong point of eDNA metabarcoding studies is that obtained reads can always be reassessed in the future when reference databases become more complete. Thus, future studies in this region may benefit from use of multiple barcode regions and databases, as well as the use of multiple taxa of sponge samplers, to improve resolution and yield of taxonomic classification. Additionally, comparative studies of nsDNA and eDNA from water filtration in the Mar Menor could improve the robustness of species community data derived from sponge eDNA in this region in the future. Finally, increasing the time resolution of sponge eDNA across the year and through long time series in the Mar Menor could also further improve understanding of the seasonal variance in taxa detected by sponges. Importantly, this nsDNA method could also be applied to samples collected in previous surveys in the study area in order to provide a picture of the ecosystem complexity in the past.

## Conclusion

Continuous, comprehensive biodiversity monitoring in marine environments can be a costly and difficult endeavor, particularly in areas that are marked by frequent disturbances and dynamic biological communities. In this study, we employed the increasingly popular sponge nsDNA technique, using samples collected from different species of *Haliclona* to characterize the faunal communities in proximity of artificial structures in the Mar Menor lagoon, a large saltwater lagoon of biological significance that has experienced frequent disturbances over the past decades. With minimal time and financial effort, we were able to use this method to detect 76 taxa, as well as to characterise the shifts in species assemblages in two different sub-locations of the lagoon and over two seasons. Interestingly, we detected invasive and allochthonous species in Mar Menor, a highly important issue for conservation, and our approach, could provide confidence in invasive species detections and their densities. Finally, by employing phylogenetic analyses in combination with eDNA metabarcoding, we were also able to provide further evidence of the taxa detections we obtained. Our findings further support the utility of the sponge nsDNA technique for continuous and comprehensive biodiversity monitoring in marine waters.

## Acknowledgments

APR acknowledges funding from “Monitoring and predictive analysis of the ecological state evolution of the Mar Menor lagoon ecosystem and prevention of impacts 2020-2022 and 2023” financed by the General Directorate of the Mar Menor of the Region of Murcia that also have contributed to the contracts of BMC and MVR. AR acknowledges funding from the SponBIODIV project (granted to AR and ST), a 2021-2022 BiodivProtect joint call for research proposals, under the Biodiversa+ Partnership co-funded by the European Commission, and with the funding organisations ‘Fundación Biodiversidad’ and FORMAS, the grant NE/T007028/1 from the UK Natural Environment Research Council (to SM and AR), an intramural grant from CSIC (PIE-202030E006) to AR, and three grants from the Spanish Ministry of Science and Innovation (RYC2018-024247-I, PID2019-105769GB-I00, and CNS2023-144571) in the framework of MCIN/AEI/10.13039/50110001103 and EI “FSE invierte en tu futuro”, to AR. ST received funding from the grants PID2020-117115GA-100 and by CNS2023-144572 funded by MCIN/AEI/10.13039/501100011033 and by the Ramón y Cajal grant RYC2021-03152-I, funded by the MCIN/AEI/10.13039/501100011033 and the European Union «NextGenerationEU/PRTR».

## Supplementary Table captions

**Supplementary Table 1.** Final ASV table for the metazoan taxa found in the study including details for number of reads, proportion by sample, and all metadata associated.

**Supplementary Table 2.** ANOVA tests for differences on beta diversity of each region, site, and the combination of both, and PERMANOVA results using Bray-Curtis distances.

