## Supplementary Figures for "Spatial and seasonal biodiversity variation in a large Mediterranean lagoon using environmental DNA metabarcoding through sponge tissue collection"

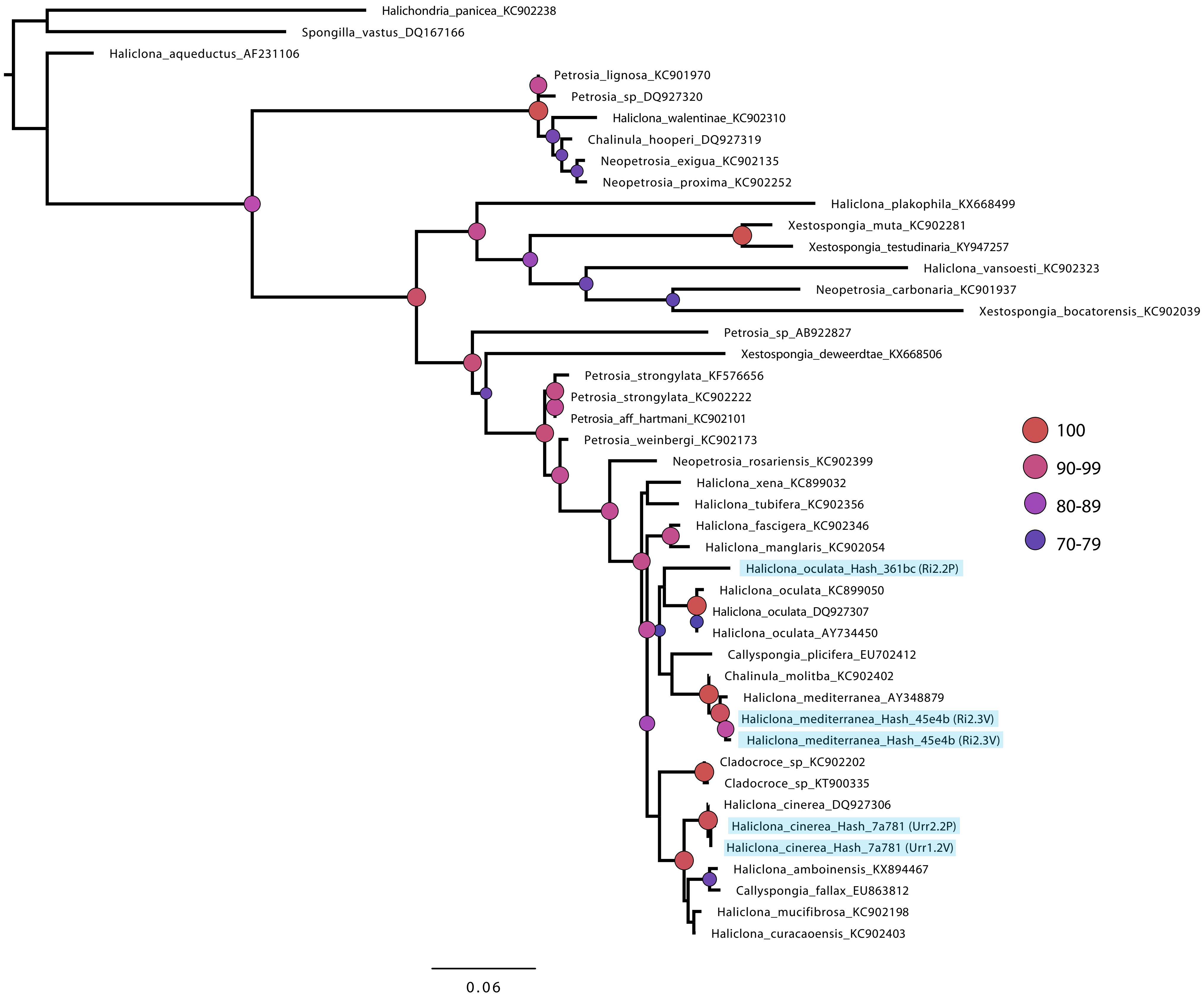
Halichondria_panicea_KC902238
Spongilla_vastus_DQ167166
Haliclona_aqueductus_AF231106
Petrosia_lignosa_KC901970
Petrosia_sp_DQ927320
Haliclona_walentinae_KC902310
Chalinula_hooperi_DQ927319
Neopetrosia_exigua_KC902135
Neopetrosia_proxima_KC902252
Haliclona_plakophila_KX668499
Xestospongia_muta_KC902281
Xestospongia_testudinaria_KY947257
Haliclona_vansoesti_KC902323
Neopetrosia_carbonaria_KC901937
Xestospongia_bocatorensis_KC902039
Petrosia_sp_AB922827
Xestospongia_deweerdtae_KX668506
Petrosia_strongylata_KF576656
Petrosia_strongylata_KC902222
Petrosia_aff_hartmani_KC902101
Petrosia_weinbergi_KC902173
Neopetrosia_rosariensis_KC902399
Haliclona_xena_KC899032
Haliclona_tubifera_KC902356
Haliclona_fascigera_KC902346
Haliclona_manglaris_KC902054
Haliclona_oculata_Hash_361bc (Ri2.2P)
Haliclona_oculata_KC899050
Haliclona_oculata_DQ927307
Haliclona_oculata_AY734450
Callyspongia_plicifera_EU702412
Chalinula_molitba_KC902402
Haliclona_mediterranea_AY348879
Haliclona_mediterranea_Hash_45e4b (Ri2.3V)
Haliclona_mediterranea_Hash_45e4b (Ri2.3V)
Cladocroce_sp_KC902202
Cladocroce_sp_KT900335
Haliclona_cinerea_DQ927306
Haliclona_cinerea_Hash_7a781 (Urr2.2P)
Haliclona_cinerea_Hash_7a781 (Urr1.2V)
Haliclona_amboinensis_KX894467
Callyspongia_fallax_EU863812
Haliclona_mucifibrosa_KC902198
Haliclona_curacaoensis_KC902403
100
90-99
80-89
70-79
0.06

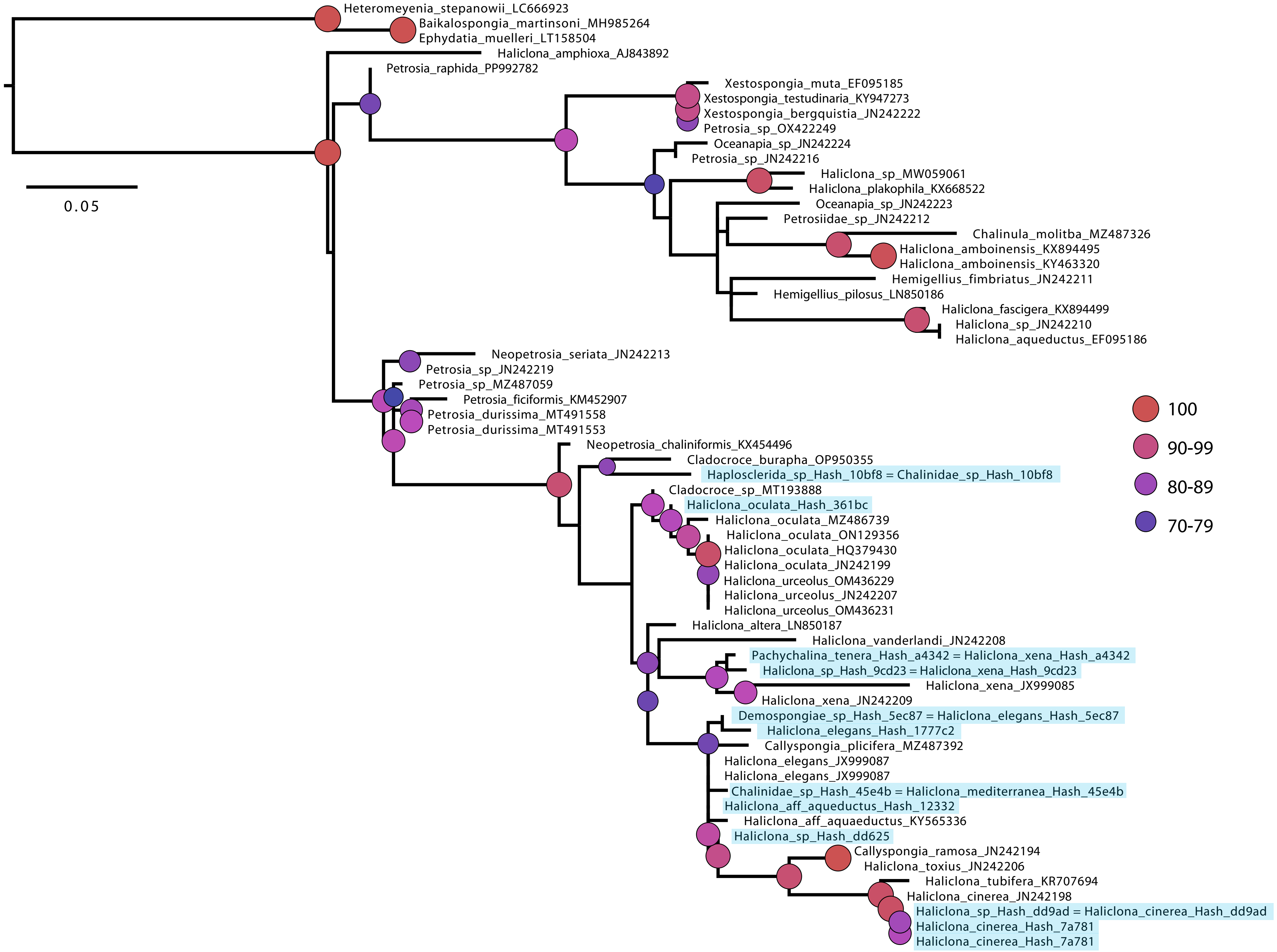

Heteromeyenia_stepanowii_LC666923
Baikalospongia_martinsoni_MH985264
Ephydatia_muelleri_LT158504
Haliclona_amphioxa_AJ843892
Petrosia_raphida_PP992782
Xestospongia_muta_EF095185
Xestospongia_testudinaria_KY947273
Xestospongia_bergquistia_JN242222
Petrosia_sp_OX422249
Oceanapia_sp_JN242224
Petrosia_sp_JN242216
Haliclona_sp_MW059061
Haliclona_plakophila_KX668522
Oceanapia_sp_JN242223
Petrosiidae_sp_JN242212
Chalinula_molitba_MZ487326
Haliclona_amboinensis_KX894495
Haliclona_amboinensis_KY463320
Hemigellius_fimbriatus_JN242211
Hemigellius_pilosus_LN850186
Haliclona_fascigera_KX894499
Haliclona_sp_JN242210
Haliclona_aqueductus_EF095186
0.05
Neopetrosia_seriata_JN242213
Petrosia_sp_JN242219
Petrosia_sp_MZ487059
Petrosia_ficiformis_KM452907
Petrosia_durissima_MT491558
Petrosia_durissima_MT491553
Neopetrosia_chaliniformis_KX454496
Cladocroce_burapha_OP950355
Haplosclerida_sp_Hash_10bf8 = Chalinidae_sp_Hash_10bf8
Cladocroce_sp_MT193888
Haliclona_oculata_Hash_361bc
Haliclona_oculata_MZ486739
Haliclona_oculata_ON129356
Haliclona_oculata_HQ379430
Haliclona_oculata_JN242199
Haliclona_urceolus_OM436229
Haliclona_urceolus_JN242207
Haliclona_urceolus_OM436231
Haliclona_altera_LN850187
Haliclona_vanderlandi_JN242208
Pachychalina_tenera_Hash_a4342 = Haliclona_xena_Hash_a4342
Haliclona_sp_Hash_9cd23 = Haliclona_xena_Hash_9cd23
Haliclona_xena_JX999085
Haliclona_xena_JN242209
Demospongiae_sp_Hash_5ec87 = Haliclona_elegans_Hash_5ec87
Haliclona_elegans_Hash_1777c2
Callyspongia_plicifera_MZ487392
Haliclona_elegans_JX999087
Haliclona_elegans_JX999087
Chalinidae_sp_Hash_45e4b = Haliclona_mediterranea_Hash_45e4b
Haliclona_aff_aqueductus_Hash_12332
Haliclona_aff_aquaeductus_KY565336
Haliclona_sp_Hash_dd625
Callyspongia_ramosa_JN242194
Haliclona_toxius_JN242206
Haliclona_tubifera_KR707694
Haliclona_cinerea_JN242198
Haliclona_sp_Hash_dd9ad = Haliclona_cinerea_Hash_dd9ad
Haliclona_cinerea_Hash_7a781
Haliclona_cinerea_Hash_7a781
100
90-99
80-89
70-79

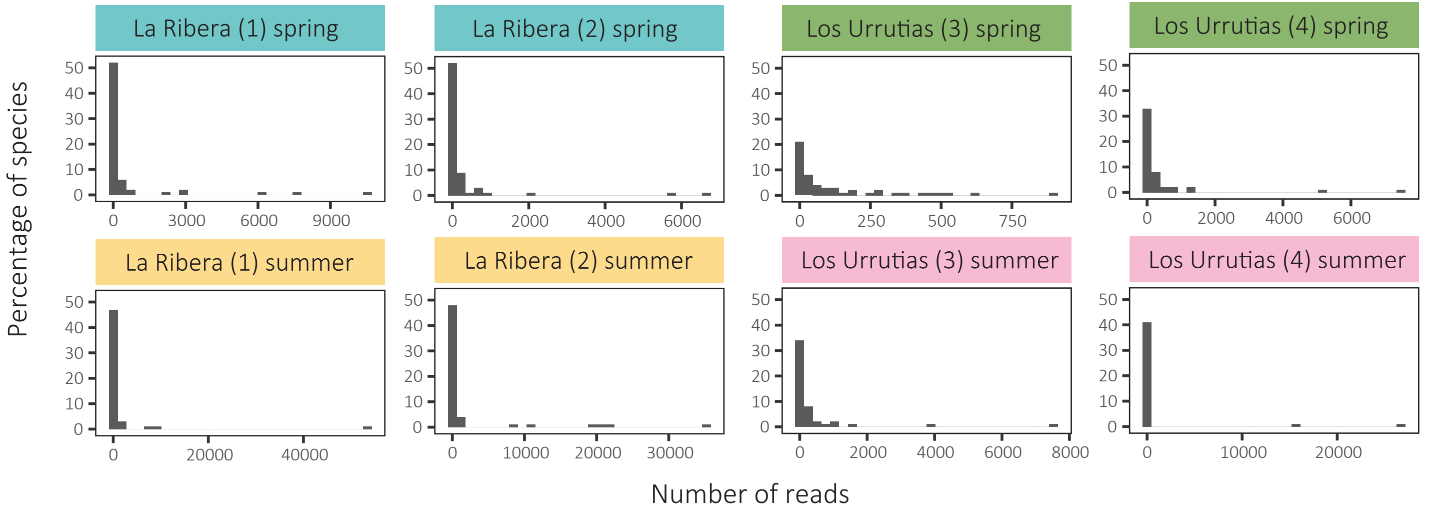
La Ribera (1) spring
La Ribera (2) spring
Los Urrutias (3) spring
Los Urrutias (4) spring
La Ribera (1) summer
La Ribera (2) summer
Los Urrutias (3) summer
Los Urrutias (4) summer
Percentage of species
Number of reads
0
10
20
30
40
50
0
3000
6000
9000
2000
4000
6000
250
500
750
20000
40000
10000
30000
8000

Rarefaction/Extrapolation Curves

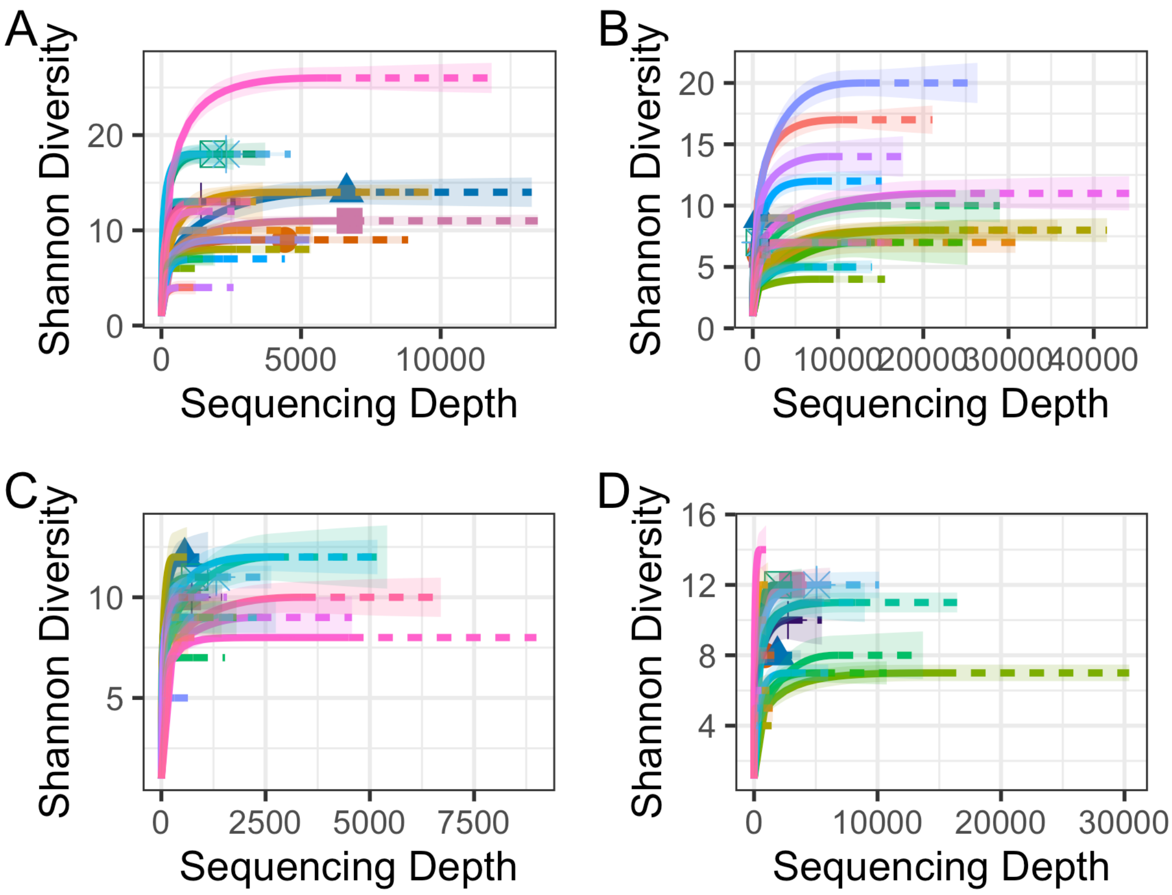


Supplementary Figure X: Rarefaction and extrapolation curves for all PCR replicates from the Mar Menor sponge samples in: la Ribera in the spring (A) and summer (B), and Los Urrutias in the spring (A) and summer (B); bootstrapping = 100.

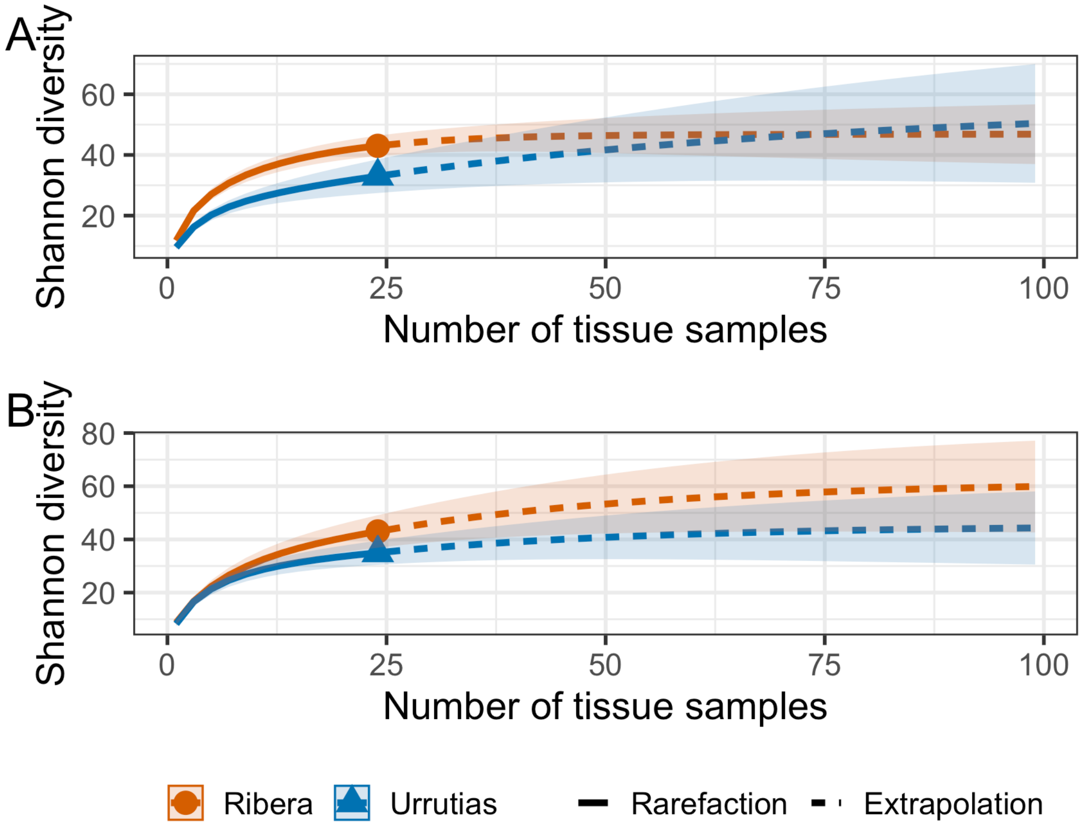


Supplementary Figure X. Rarefaction and extrapolation curves of sampling replicates in the respective sampling areas of the Mar Menor in the (A) spring and (B) summer; bootstrapping = 100.

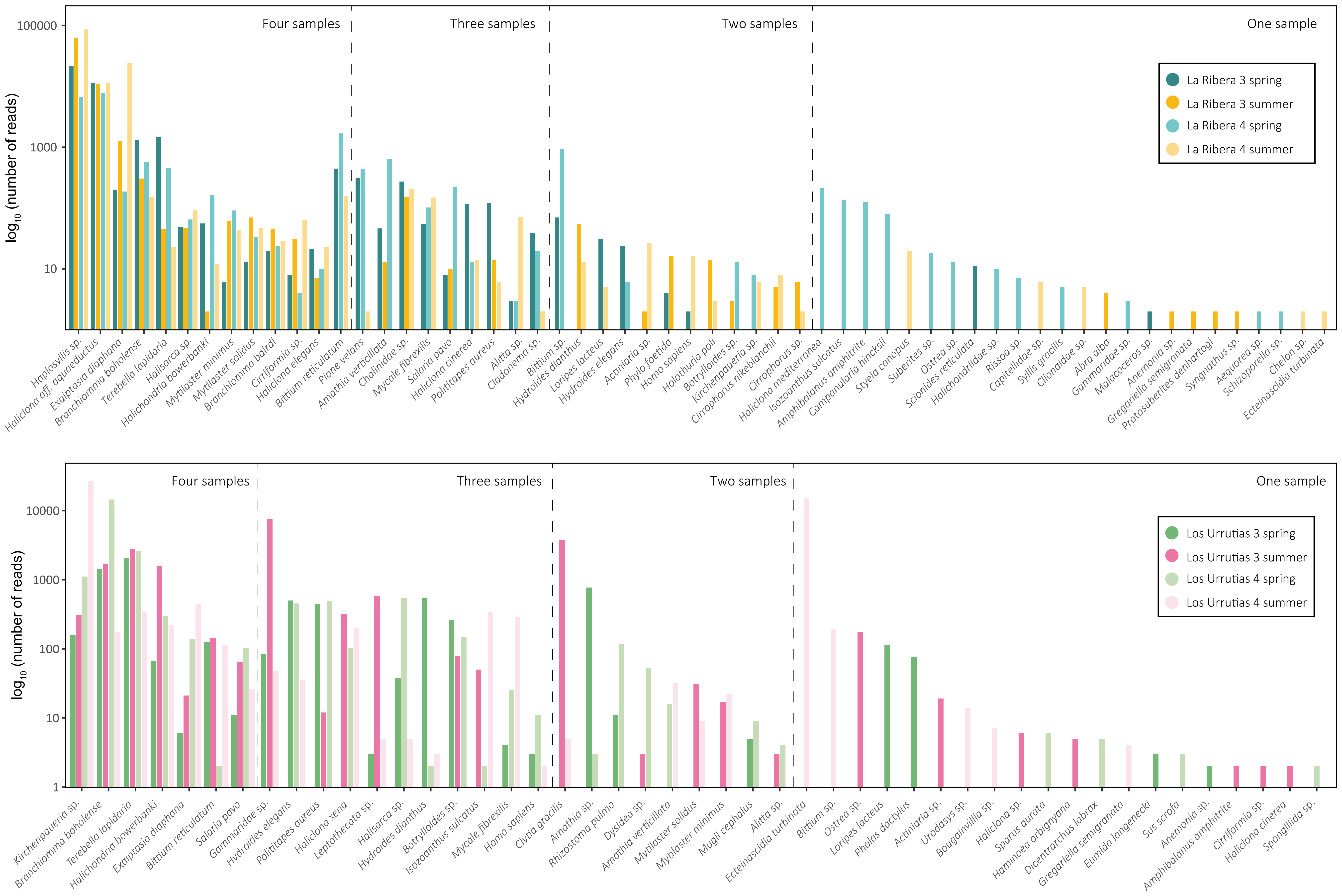

log10 (number of reads)
100000
1000
10
Four samples
Three samples
Two samples
One sample
La Ribera 3 spring
La Ribera 3 summer
La Ribera 4 spring
La Ribera 4 summer
Haplosyllis sp.
Haliclona aff. aqueductus
Exaiptasia diaphana
Branchiomma boholense
Terebella lapidaria
Halisarca sp.
Halichondria bowerbanki
Mytilaster minimus
Mytilaster solidus
Branchiomma bairdi
Cirriformia sp.
Haliclona elegans
Bittium reticulatum
Pione velans
Amathia verticillata
Chalinidae sp.
Mycale fibrexilis
Salaria pavo
Haliclona cinerea
Polititapes aureus
Alitta sp.
Cladonema sp.
Bittium sp.
Hydroides dianthus
Loripes lacteus
Hydroides elegans
Actiniaria sp.
Phylo foetida
Homo sapiens
Holothuria poli
Botrylloides sp.
Kirchenpaueria sp.
Cirrophorus nikebianchii
Cirrophorus sp.
Haliclona mediterranea
Isozoanthus sulcatus
Amphibalanus amphitrite
Campanularia hincksii
Styela canopus
Suberites sp.
Ostrea sp.
Scionides reticulata
Halichondriidae sp.
Rissoa sp.
Capitellidae sp.
Syllis gracilis
Clionaidae sp.
Abra alba
Gammaridae sp.
Malacoceros sp.
Anemonia sp.
Gregariella semigranata
Protosuberites denhartogi
Syngnathus sp.
Aequorea sp.
Schizoporella sp.
Chelon sp.
Ecteinascidia turbinata
log10 (number of reads)
10000
1000
100
10
1
Four samples
Three samples
Two samples
One sample
Los Urrutias 3 spring
Los Urrutias 3 summer
Los Urrutias 4 spring
Los Urrutias 4 summer
Kirchenpaueria sp.
Branchiomma boholense
Terebella lapidaria
Halichondria bowerbanki
Exaiptasia diaphana
Bittium reticulatum
Salaria pavo
Gammaridae sp.
Hydroides elegans
Polititapes aureus
Haliclona xena
Leptothecata sp.
Halisarca sp.
Hydroides dianthus
Botrylloides sp.
Isozoanthus sulcatus
Mycale fibrexilis
Homo sapiens
Clytia gracilis
Amathia sp.
Rhizostoma pulmo
Dysidea sp.
Amathia verticillata
Mytilaster solidus
Mytilaster minimus
Mugil cephalus
Alitta sp.
Ecteinascidia turbinata
Bittium sp.
Ostrea sp.
Loripes lacteus
Pholas dactylus
Actiniaria sp.
Urodasys sp.
Bougainvillia sp.
Haliclona sp.
Sparus aurata
Haminoea orbignyana
Dicentrarchus labrax
Gregariella semigranata
Eumida langenecki
Sus scrofa
Anemonia sp.
Amphibalanus amphitrite
Cirriformia sp.
Haliclona cinerea
Spongillida sp.

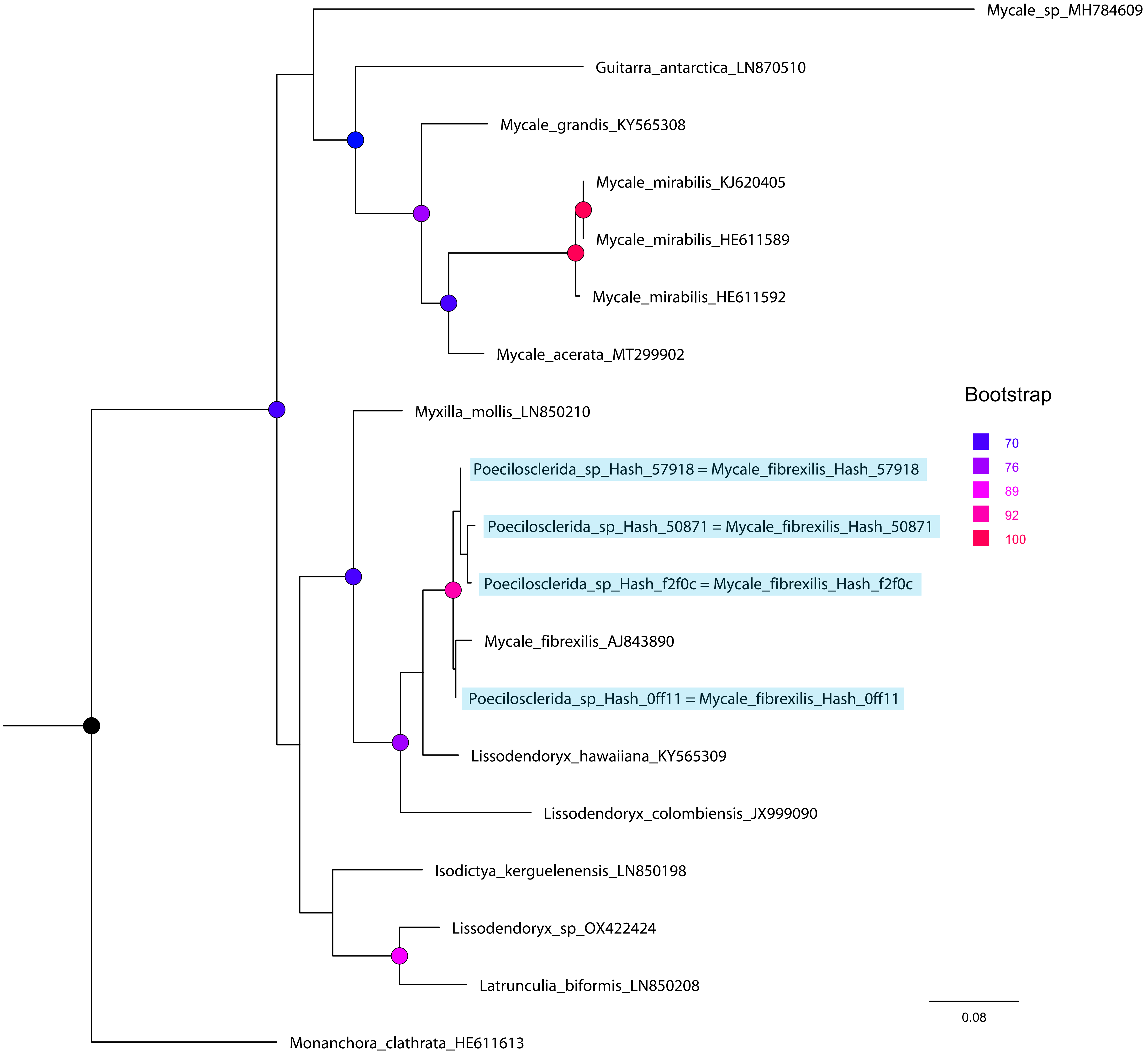
Mycale_sp_MH784609
Guitarra_antarctica_LN870510
Mycale_grandis_KY565308
Mycale_mirabilis_KJ620405
Mycale_mirabilis_HE611589
Mycale_mirabilis_HE611592
Mycale_acerata_MT299902
Myxilla_mollis_LN850210
Poecilosclerida_sp_Hash_57918 = Mycale_fibrexilis_Hash_57918
Poecilosclerida_sp_Hash_50871 = Mycale_fibrexilis_Hash_50871
Poecilosclerida_sp_Hash_f2f0c = Mycale_fibrexilis_Hash_f2f0c
Mycale_fibrexilis_AJ843890
Poecilosclerida_sp_Hash_0ff11 = Mycale_fibrexilis_Hash_0ff11
Lissodendoryx_hawaiiana_KY565309
Lissodendoryx_colombiensis_JX999090
Isodictya_kerguelenensis_LN850198
Lissodendoryx_sp_OX422424
Latrunculia_biformis_LN850208
Monanchora_clathrata_HE611613
Bootstrap
70
76
89
92
100
0.08

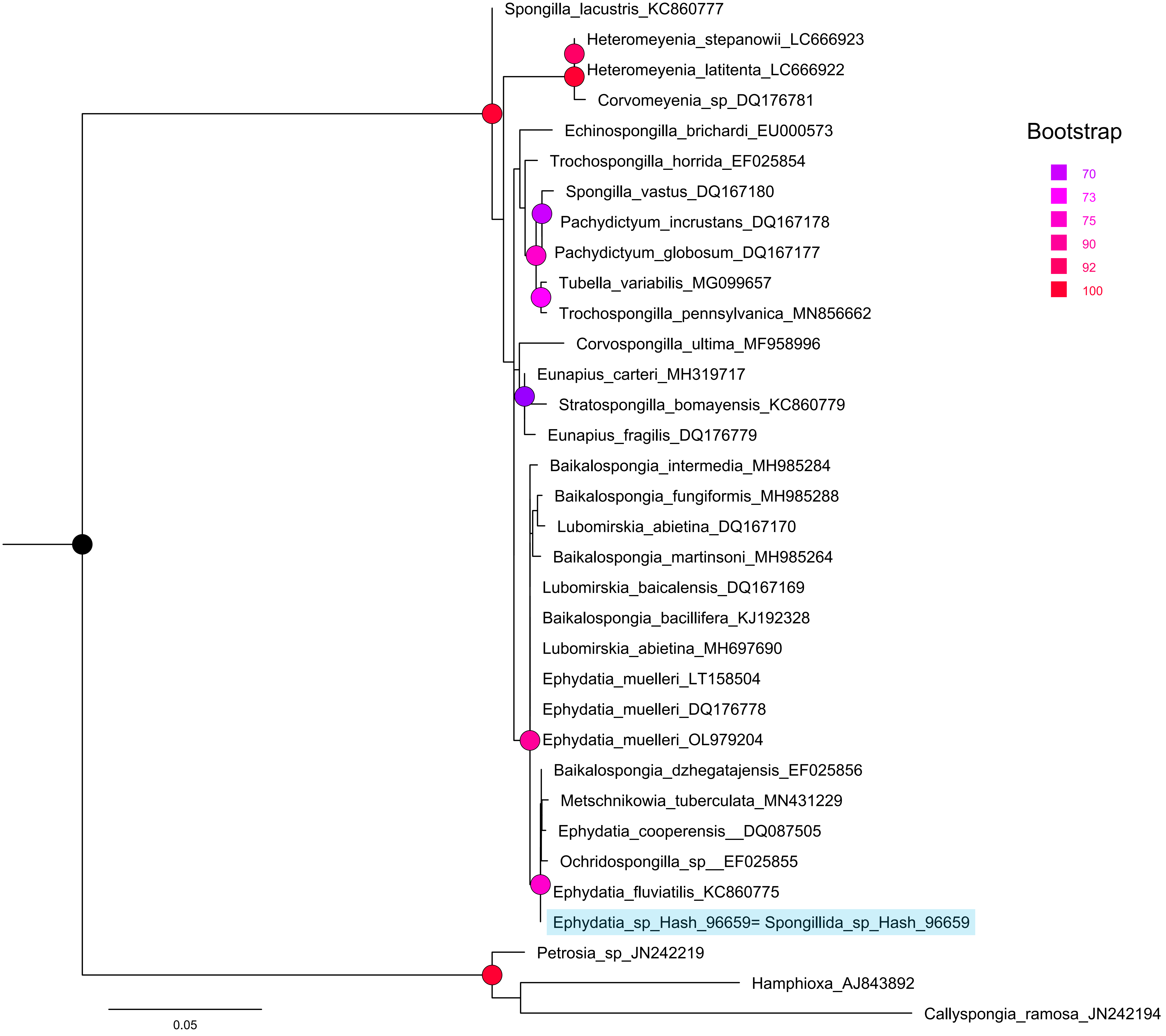

Spongilla_lacustris_KC860777
Heteromeyenia_stepanowii_LC666923
Heteromeyenia_latitenta_LC666922
Corvomeyenia_sp_DQ176781
Echinospongilla_brichardi_EU000573
Trochospongilla_horrida_EF025854
Spongilla_vastus_DQ167180
Pachydictyum_incrustans_DQ167178
Pachydictyum_globosum_DQ167177
Tubella_variabilis_MG099657
Trochospongilla_pennsylvanica_MN856662
Corvospongilla_ultima_MF958996
Eunapius_carteri_MH319717
Stratospongilla_bomayensis_KC860779
Eunapius_fragilis_DQ176779
Baikalospongia_intermedia_MH985284
Baikalospongia_fungiformis_MH985288
Lubomirskia_abietina_DQ167170
Baikalospongia_martinsoni_MH985264
Lubomirskia_baicalensis_DQ167169
Baikalospongia_bacillifera_KJ192328
Lubomirskia_abietina_MH697690
Ephydatia_muelleri_LT158504
Ephydatia_muelleri_DQ176778
Ephydatia_muelleri_OL979204
Baikalospongia_dzhegatajensis_EF025856
Metschnikowia_tuberculata_MN431229
Ephydatia_cooperensis__DQ087505
Ochridospongilla_sp__EF025855
Ephydatia_fluviatilis_KC860775
Ephydatia_sp_Hash_96659= Spongillida_sp_Hash_96659
Petrosia_sp_JN242219
Hamphioxa_AJ843892
Callyspongia_ramosa_JN242194
0.05
Bootstrap
70
73
75
90
92
100

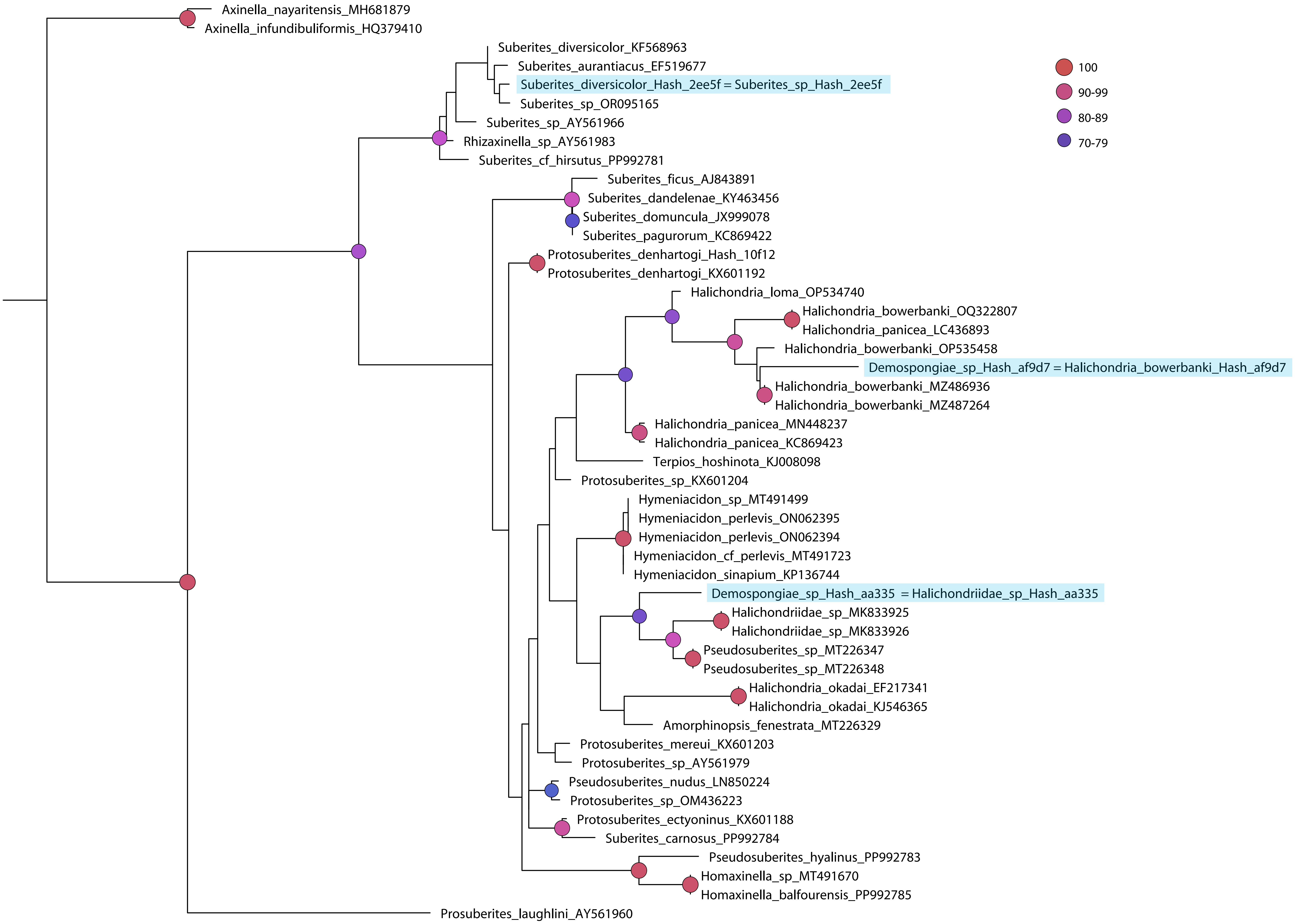

Axinella_nayaritensis_MH681879
Axinella_infundibuliformis_HQ379410
Suberites_diversicolor_KF568963
Suberites_aurantiacus_EF519677
Suberites_diversicolor_Hash_2ee5f = Suberites_sp_Hash_2ee5f
Suberites_sp_OR095165
Suberites_sp_AY561966
Rhizaxinella_sp_AY561983
Suberites_cf_hirsutus_PP992781
Suberites_ficus_AJ843891
Suberites_dandelenae_KY463456
Suberites_domuncula_JX999078
Suberites_pagurorum_KC869422
Protosuberites_denhartogi_Hash_10f12
Protosuberites_denhartogi_KX601192
Halichondria_loma_OP534740
Halichondria_bowerbanki_OQ322807
Halichondria_panicea_LC436893
Halichondria_bowerbanki_OP535458
Demospongiae_sp_Hash_af9d7 = Halichondria_bowerbanki_Hash_af9d7
Halichondria_bowerbanki_MZ486936
Halichondria_bowerbanki_MZ487264
Halichondria_panicea_MN448237
Halichondria_panicea_KC869423
Terpios_hoshinota_KJ008098
Protosuberites_sp_KX601204
Hymeniacidon_sp_MT491499
Hymeniacidon_perlevis_ON062395
Hymeniacidon_perlevis_ON062394
Hymeniacidon_cf_perlevis_MT491723
Hymeniacidon_sinapium_KP136744
Demospongiae_sp_Hash_aa335 = Halichondriidae_sp_Hash_aa335
Halichondriidae_sp_MK833925
Halichondriidae_sp_MK833926
Pseudosuberites_sp_MT226347
Pseudosuberites_sp_MT226348
Halichondria_okadai_EF217341
Halichondria_okadai_KJ546365
Amorphinopsis_fenestrata_MT226329
Protosuberites_mereui_KX601203
Protosuberites_sp_AY561979
Pseudosuberites_nudus_LN850224
Protosuberites_sp_OM436223
Protosuberites_ectyoninus_KX601188
Suberites_carnosus_PP992784
Pseudosuberites_hyalinus_PP992783
Homaxinella_sp_MT491670
Homaxinella_balfourensis_PP992785
Prosuberites_laughlini_AY561960
100
90-99
80-89
70-79

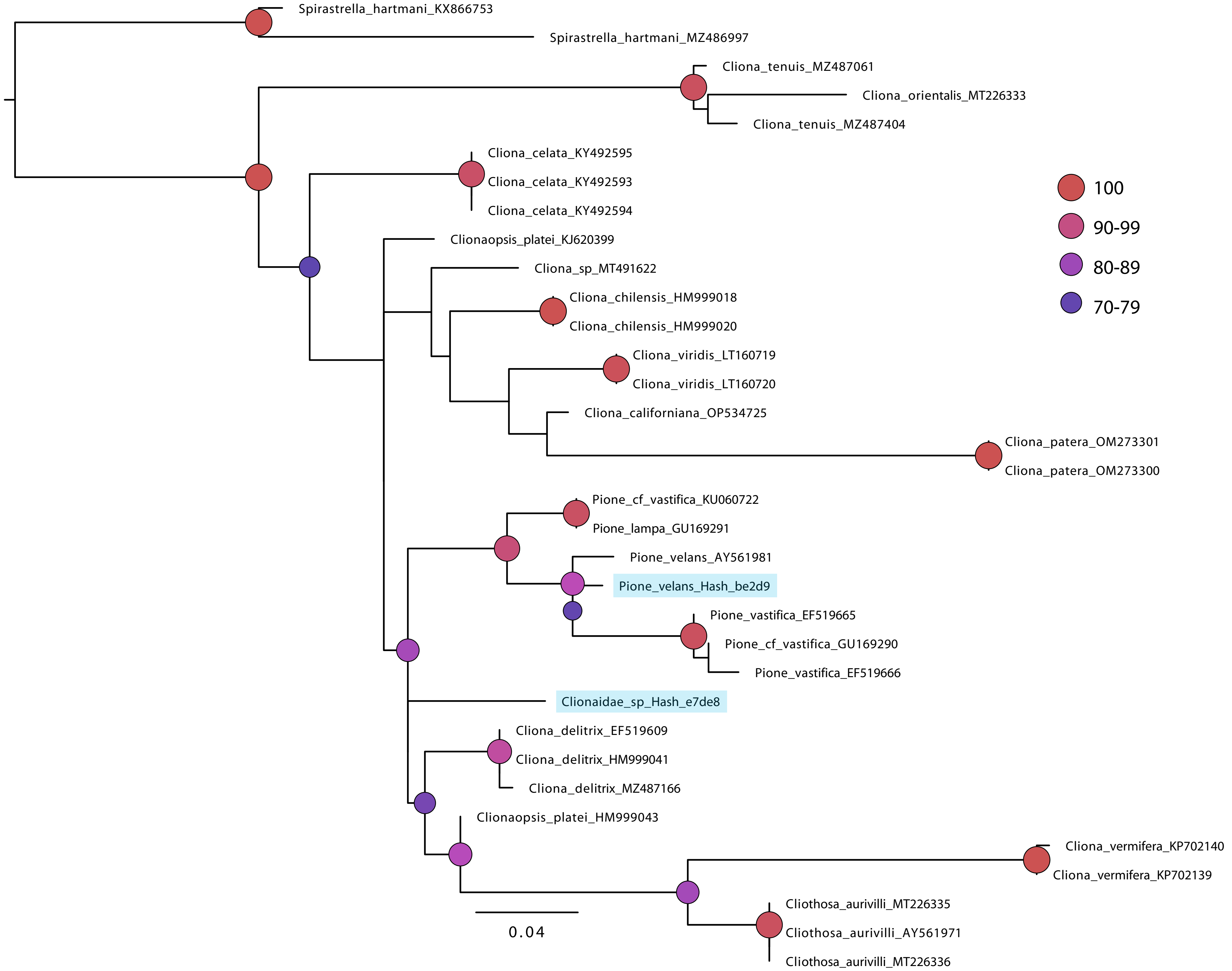
Spirastrella_hartmani_KX866753
Spirastrella_hartmani_MZ486997
Cliona_tenuis_MZ487061
Cliona_orientalis_MT226333
Cliona_tenuis_MZ487404
Cliona_celata_KY492595
Cliona_celata_KY492593
Cliona_celata_KY492594
Clionaopsis_platei_KJ620399
Cliona_sp_MT491622
Cliona_chilensis_HM999018
Cliona_chilensis_HM999020
Cliona_viridis_LT160719
Cliona_viridis_LT160720
Cliona_californiana_OP534725
Cliona_patera_OM273301
Cliona_patera_OM273300
Pione_cf_vastifica_KU060722
Pione_lampa_GU169291
Pione_velans_AY561981
Pione_velans_Hash_be2d9
Pione_vastifica_EF519665
Pione_cf_vastifica_GU169290
Pione_vastifica_EF519666
Clionaidae_sp_Hash_e7de8
Cliona_delitrix_EF519609
Cliona_delitrix_HM999041
Cliona_delitrix_MZ487166
Clionaopsis_platei_HM999043
Cliona_vermifera_KP702140
Cliona_vermifera_KP702139
Cliothosa_aurivilli_MT226335
Cliothosa_aurivilli_AY561971
Cliothosa_aurivilli_MT226336
0.04
100
90-99
80-89
70-79
